# ATP-independent unfolding of ubiquitin by Ufd1 initiates Cdc48/p97-mediated substrate processing

**DOI:** 10.64898/2026.03.07.710261

**Authors:** Yanzhu Wang, Zhimin Zhang, Wei He, Pengyu Wang, Zhejian Ji

## Abstract

The Cdc48 ATPase (p97 or VCP in mammals) cooperates with its cofactors Ufd1 and Npl4 to extract polyubiquitinated proteins from membranes or multi-subunit complexes, promoting their proteasomal degradation. A ubiquitin molecule in the chain is unfolded in an ATP-independent manner and initiates substrate translocation through the central pore of the ATPase. How ubiquitin is unfolded remains unclear. Here, we demonstrate that the UT3 domain of Ufd1 interacts simultaneously with two Lys48-linked ubiquitins and unfolds one of the ubiquitins by binding its C-termianl β-strand into a cleft. This unfolded ubiquitin is subsequently captured by Npl4 and Cdc48/p97. Experiments *in vitro* and in cells show that unfolding-defective mutants of Ufd1 indeed compromise Cdc48/p97 function. Our results explain how simple protein-protein interactions cause the unfolding of the remarkably stable ubiquitin. We also establish a linkage-specific recognition that involves ubiquitin unfolding.

## INTRODUCTION

Cdc48 and its mammalian homolog p97 or VCP belong to the AAA ATPase family (ATPases Associated with diverse cellular Activities). The hexameric Cdc48/p97 plays a critical role in various cellular processes. In one of the best studied pathways, the ATPase acts in concert with a heterodimeric cofactor, consisting of Ufd1 and Npl4^1–4^.

This Cdc48/p97–Ufd1–Npl4 ATPase complex unfolds protein substrates carrying a Lys48-linked polyubiquitin chain and extracts them from membranes or macromolecules, prior to their degradation by the proteasome^2,5–7^. Many cancer cells often upregulate the p97 ATPase complex in order to bolster protein turnover. Thus, targeting human p97 or its cofactors has emerged to be a new strategy to inhibit protein degradation and to intervene cancer development^8–10^.

Although both the Cdc48/p97 ATPase complex and the proteasome recruit polyubiquitinated substrates, they employ distinct mechanisms. The proteasome captures an unstructured region from the substrate, which is often longer than 20 amino acids^11,12^. This unstructured segment is subsequently used as the initiator for polypeptide translocation and degradation^13,14^. In contrast, the Cdc48/p97 ATPase complex does not require such an unstructured segment from its substrate. Instead, it captures a ubiquitin molecule in unfolded state, forming the initiation complex (**Extended Data Fig. 1a**)^15,16^. This unfolded ubiquitin enhances polyubiquitin binding to the ATPase complex and serves as the initiator segment for substrate processing^17^.

Strikingly, ATP hydrolysis is not required for the unfolding of initiator ubiquitin and the assembly of the initiation complex^15^. Because ubiquitin is generally perceived as a well-folded protein with superior thermostability, the mechanism by which the initiator ubiquitin is unfolded is interesting but remains mysterious.

One possible hypothesis is that ubiquitin binding with a domain within the ATPase complex induces ubiquitin unfolding. Earlier cryogenic electron microscopy (cryo-EM) structures of the initiation complex have revealed that Cdc48 central pore and Npl4 groove interact with the N-terminal and middle segment of the initiator ubiquitin, respectively^15^. These two interactions, however, are likely the consequence rather than the cause of ubiquitin unfolding. In the folded state, ubiquitin adopts a β-grasp fold with an alpha helix, five β strands (β1-β5), and two short 3_10_ helices (**Extended Data Fig. 1b,c**)^18^. The N-terminal β1 interacts with β2 and the C-terminal β5, forming a β-sheet. The β1-β5 interaction has to be disrupted before the N-terminus of ubiquitin can be inserted into the ATPase central pore. Similarly, the hydrophobic surface of ubiquitin alpha helix that is normally packed inside and not exposed in the folded state binds to the Npl4 groove in the initiation complex, indicating that deformation of the β-grasp fold is a prerequisite for ubiquitin to bind with Npl4. These arguments imply that another ubiquitin-binding domain, other than Npl4 and Cdc48, might be responsible for triggering ubiquitin unfolding.

Within the ATPase complex, the UT3 domain of Ufd1 is another known ubiquitin-binding domain. Ye and colleagues first reported that UT3 of *Saccharomyces cerevisiae* Ufd1 is not only required but also sufficient to bind with Lys48-linked polyubiquitin^19^. Later, the structure of UT3 was determined using nuclear magnetic resonance (NMR), revealing two subdomains – a double-psi, β-barrel-fold UT3n and a mixed-α/β-roll-fold UT3c^20^.

Evolutionary conservation analysis identified two highly conserved surfaces on UT3 (**Extended Data Fig. 1d,e**). One is a cavity at the ridge surface of UT3n, encircled by α2, the α2-β3 linker, and β4. The other is a cleft within UT3c, sandwiched by α4, the β11-β12 linker, and β12. Hereafter, these two conserved sites on UT3n and UT3c will be referred as the ridge and cleft site, respectively. Previous efforts to map polyubiquitin-binding sites on Ufd1 using the hydrogen-deuterium exchange coupled with mass spectrometry technique (HDX/MS) found that both the ridge and cleft sites showed reduced HDX rate upon polyubiquitin binding (**Extended Data Fig. 1f**)^15^, indicating these regions might directly interact with ubiquitin. Moreover, ubiquitin binding to UT3 ridge was further characterized by a combination of AlphaFold modeling, mutagenesis, and fluorescence resonance energy transfer (FRET) experiments, which identified Ufd1 Leu55 and ubiquitin Ile44 as the critical residues^21^. Despite of these results, it remains to be tested whether UT3 induces ubiquitin unfolding.

Ufd1 is indispensable for the polyubiquitinated substrate processing by the Cdc48/p97 ATPase complex. Yet the exact role of Ufd1 remains mysterious. Because the UT3 domain binds to polyubiquitin, Ufd1 has been postulated to simply bridge the interaction between polyubiquitin and the Cdc48/p97 ATPase. However, the UT3-ubiquitin binding appeared to be quite weak, comparing with Npl4 and other canonical ubiquitin-binding proteins^22^. How such weak binding contributes significantly to the ATPase complex function remains to be elucidated.

Herein, we demonstrate that the UT3 domain of Ufd1 is required and sufficient to induce ubiquitin unfolding in an ATP-independent manner. Through its conserved ridge and cleft sites, UT3 simultaneously interact with two ubiquitin molecules within a polyubiquitin chain. In particular, the UT3 cleft binding with ubiquitin C-terminal β5 strand causes conformational alteration of the ubiquitin, eventually leads to its unfolding. This unique ubiquitin-unfolding function of Ufd1 is crucial for substrate processing by the Cdc48/p97 ATPase complex both *in vitro* and *in vivo*. Our work uncovers an unconventional function of the Ufd1 cofactor and clarifies how the initiator ubiquitin is unfolded. It further suggests that simple protein-protein interactions can trigger the unfolding of ubiquitin molecules with remarkable stability.

## RESULTS

### The UT3 domain of Ufd1 induces ubiquitin unfolding

To directly examine whether UT3 triggers ubiquitin unfolding, we adapted an unfolding reporter of ubiquitin and assayed its folding status via dye accessibility^17^. To this end, the Ile3 residue of ubiquitin was substituted to a cysteine (Ub^I3C^). In a folded ubiquitin, I3C side chain folds inside a hydrophobic core (**Extended Data Fig. 1b**) and is therefore not accessible to fluorescent dye-conjugated maleimide moieties (Dy-maleimide) (**Fig. 1a**). When ubiquitin is unfolded, I3C becomes exposed and can react with Dy-maleimide, resulting in covalent labeling by the fluorescent dye. Thus, fluorescence intensity on ubiquitin is proportional to the number of unfolded molecules. The I3C ubiquitin was used to generate Lys48-linked chains [Ub^I3C^(n)]. Further cation-exchange chromatography resulted in fractions that contain primarily di- or tri-ubiquitins (**Extended Data Fig. 1h**). In the dye accessibility assay, I3C di-ubiquitin pre-incubated with UT3 (residues 1-201 of Ufd1; all proteins are referred to *S. cerevisiae* proteins unless noted otherwise) showed remarkable increase of dye labeling, comparing to the one without UT3 preincubation (**Fig. 1b**; lane 3 versus 2). Notably, no ATP was added in this assay, indicating that UT3 protein can induce ubiquitin unfolding in an ATP-independent manner.

**Fig. 1:**
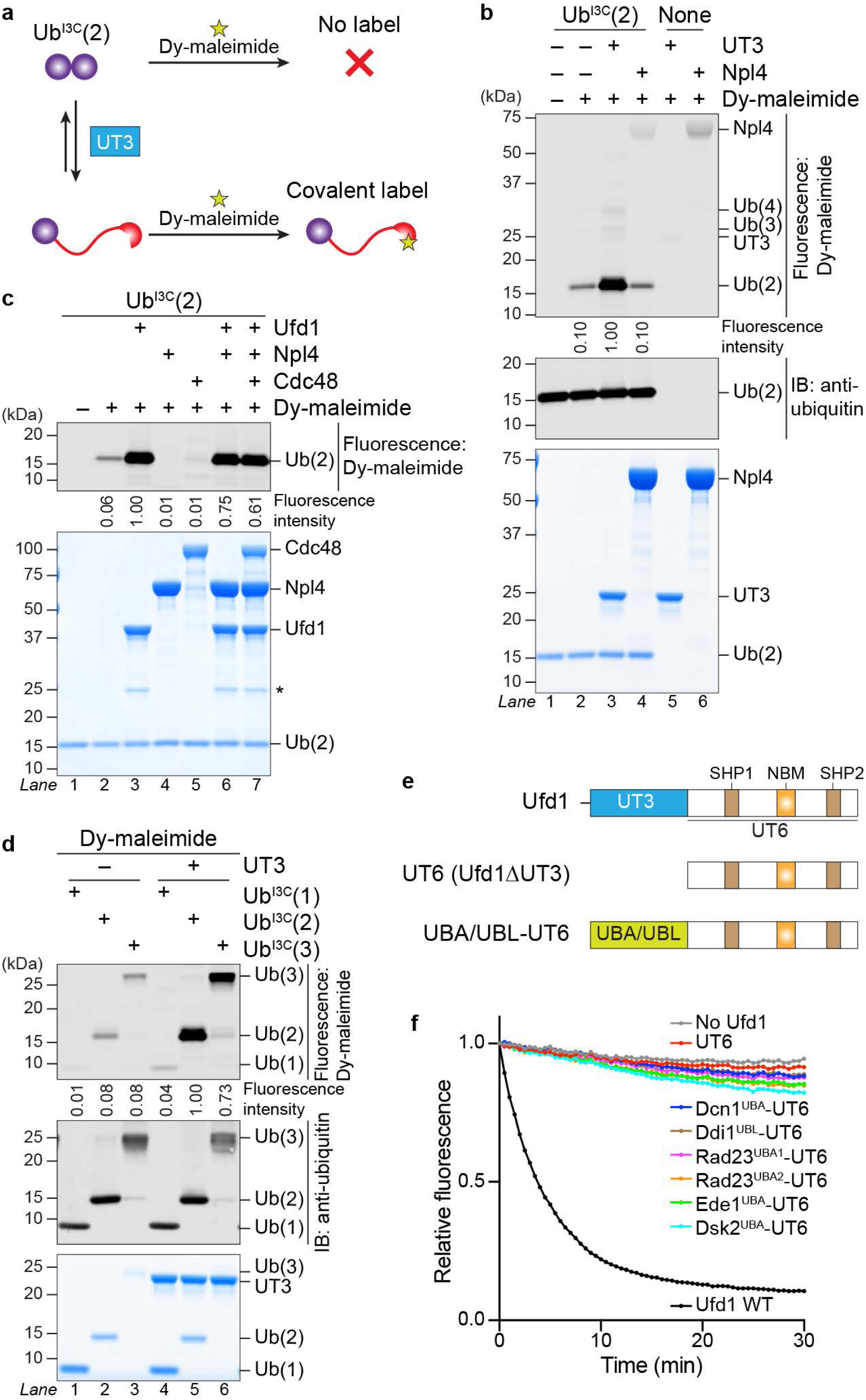
The UT3 domain of Ufd1 induces ubiquitin unfolding in an ATP-independent manner. **a,b,** Dye accessibility assay was carried out with K48-linked di-ubiquitin of the I3C mutant [Ub^I3C^(2)] that was preincubated with UT3 or Npl4. When treated with fluorescent dye-conjugated maleimide moieties (Dy-maleimide), only unfolded I3C ubiquitins were covalently labeled by the fluorescent dye (**a**). After the reaction, protein samples were analyzed by dye fluorescence scanning, immunoblotting (IB) against ubiquitin, and Coomassie blue staining (**b**). Intensities of the dye fluorescence on Ub^I3C^(2) were quantified and normalized to the strongest signal. **c,** Dye accessibility assay was conducted as in (**b**), except that the preincubations were carried out with full-length Ufd1, Npl4, and/or Cdc48. Asterisk indicates a degradation band in the purified Ufd1. **d,** Dye accessibility assays as in (**b**) were conducted with I3C mono-, di-, and tri-ubiquitin [Ub^I3C^(1), Ub^I3C^(2), and Ub^I3C^(3), respectively]. **e,f,** Chimera Ufd1 proteins were constructed by substituting the UT3 domain with ubiquitin-binding UBA or UBL domains (**e**). Each of these chimeras was incubated with polyubiquitinated, photo-converted Dendra substrate as well as Npl4 and Cdc48. Upon addition of ATP, the red fluorescence of photo-converted Dendra was followed over a time course of 30 min (**f**). Intensities of Dendra fluorescence were normalized to the zero-timepoint values of each reaction and plotted against time.

Similar to UT3, full-length Ufd1 also triggered the unfolding of I3C di-ubiquitin (**Fig. 1c**). In contrast, preincubation with Npl4 or Cdc48 did not stimulate ubiquitin unfolding, despite that Npl4 interacts with polyubiquitin as well^22–24^. Interestingly, addition of Npl4 and Cdc48 to Ufd1 did not further enhance ubiquitin unfolding (**Fig. 1c**; lanes 6-7). This result is not consistent with the previous notion, where dye accessibility of I3C polyubiquitin was stimulated by a concerted action of Ufd1, Npl4, and Cdc48^17^. Lack of a cooperativity on di-ubiquitin is likely because di-ubiquitin is too short to engage with all three proteins simultaneously. Indeed, early work demonstrated that the ATPase complex only recognizes polyubiquitin chains with five or more ubiquitins^4,6^. These lines of evidence suggest that UT3 is the minimal domain to unfold ubiquitin; Npl4 or Cdc48 does not trigger ubiquitin unfolding directly, but they may stabilize the unfolded ubiquitin when the chain is long enough.

We then compared the unfolding tendency of ubiquitin chains with various lengths. While dye labeling of di- and tri-ubiquitin were largely stimulated by UT3, the labeling of I3C mono-ubiquitin appeared to be marginal even in the presence of UT3 (**Fig. 1d**). To further examine the linkage specificity of UT3, we also generated linear and Lys63-linked I3C di-ubiquitin. Dye accessibility to these two types of di-ubiquitin did not change upon the addition of UT3, as opposed to the Lys48-type (**Extended Data Fig. 2a**). This implies that UT3 has a specificity towards Lys48-linked ubiquitins.

### Conventional ubiquitin-binding domains cannot substitute UT3 for its function in the ATPase complex

To test if other ubiquitin-binding domains can be a functional replacement, we substituted the UT3 domain with the ubiquitin-association (UBA) domains of Dcn1, Ede1, Dsk2, and Rad23, as well as the ubiquitin-like (UBL) domain of Ddi1. Dsk2, Rad23, and Ddi1 are well-known shuttling factors that contain both UBA and UBL domains^25^. Dsk2 contains one UBA and Rad23 contains two (designated as UBA1 and UBA2). Polyubiquitin binding to the UBA domains of Dsk2 and Rad23 have been reconstituted *in vitro*^22^. Recent work revealed that full-length Ddi1 requires its UBL, not UBA, for polyubiquitin binding^26^. In addition, structures have been reported for ubiquitin in complex with Ede1^UBA^ (PDB: 2G3Q)^27^, Dsk2^UBA^ (PDB: 1WR1 and 4UN2)^28,29^, or Ddi1^UBL^ (PDB: 2MRO and 2MWS)^30^. We therefore fused each of these UBA/UBL domains to the N-terminus of Ufd1 UT6, generating a set of chimera proteins (**Fig. 1e**).

Polyubiquitinated model substrates containing fluorescent protein Dendra or super-folder GFP (sfGFP) were generated as previously described (**Extended Data Fig. 2b,c**)^17^. Polyubiquitinated, photo-converted Dendra was used to test the substrate unfolding activity of the ATPase complex, whereas polyubiquitinated, dye-labeled sfGFP was applied in pulldown for the initiation complex assembly, as well as in photo-crosslinking experiments to assay the generation of unfolded initiator ubiquitins.

Consistent with previous data, deleting UT3 led to defects in Cdc48-dependent Dendra substrate unfolding (**Fig. 1f**; UT6) and recruitment of polyubiquitinated substrates to the ATPase complex (**Extended Data Fig. 2d**; lane 2). The UBA/UBL chimeras could restore substrate recruitment to levels that were similar with or even better than wild-type (WT) Ufd1 (**Extended Data Fig. 2d**). However, none of the chimeras supported Dendra substrate unfolding (**Fig. 1f**). Furthermore, we examined the generation of the unfolded initiator ubiquitin by photo-crosslinking of the initiator to a photo-reactive probe, benzoyl-L-phenylalanine (Bpa), at Cdc48 D1 pore loop (D324Bpa). In the presence of the UBA/UBL chimeras, few crosslinks between polyubiquitinated substrates and Cdc48 D324Bpa were observed (**Extended Data Fig. 2e**), suggesting that these UBA/UBL domains are incompetent in generating unfolded ubiquitin. Therefore, simply recruiting polyubiquitin to the ATPase complex is not sufficient to initiate substrate processing. The ubiquitin-unfolding function is unique to UT3 and is indispensable for the ATPase complex.

### UT3 simultaneously interacts with two Lys48-linked ubiquitins via its ridge and cleft

To understand the molecular mechanism by which UT3 unfolds di-ubiquitin, we used AlphaFold3^31^ to predict the complex structure. Due to technical difficulty in specifying an isopeptide bond, a structural prediction of UT3 in complex with two separate ubiquitins (UT3:2Ub) was generated instead. The two ubiquitins were predicted to bind at the ridge and cleft sites of UT3, respectively (**Fig. 2a**). This UT3:2Ub model is consistent with the high conservation of ubiquitin sequence and UT3 ridge and cleft sites (**Extended Data Fig. 1d**) and also aligns well with the previous HDX/MS results (**Extended Data Fig. 1f**)^15^. Strikingly, the cleft-bound ubiquitin (Ub^Cleft^) has its C-terminal Gly76 residue in close proximity to the Lys48 residue of the ridge-bound ubiquitin (Ub^Ridge^), as if they were covalently tethered by a Lys48-linked isopeptide bond (**Fig. 2b**). Thus, the UT3:2Ub model represents the structure of UT3 in complex with a Lys48-linked di-ubiquitin. It provides a good explanation at the molecular level why UT3 acts only on Lys48-linked ubiquitins (**Extended Data Fig. 2a**).

**Fig. 2:**
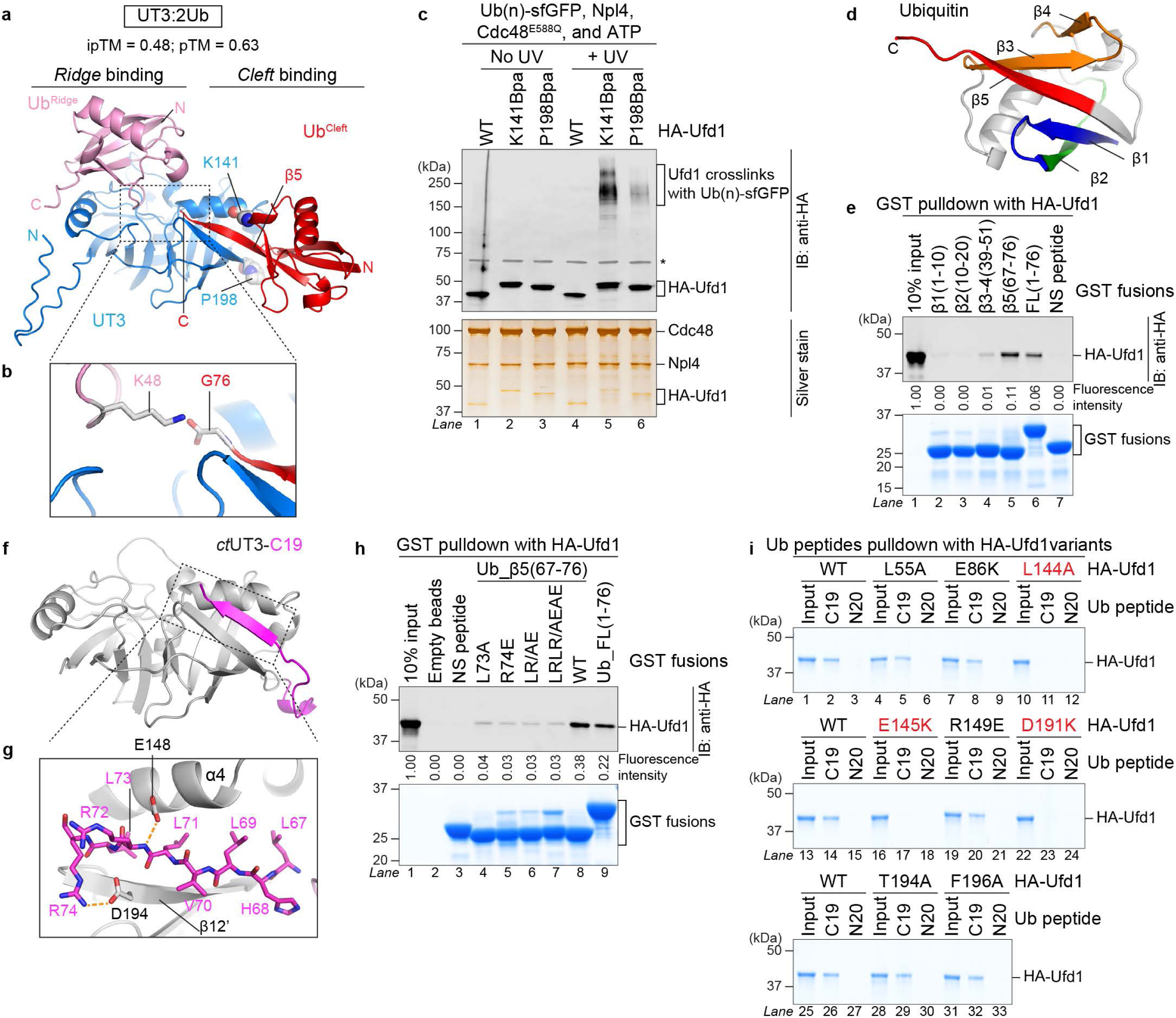
The conserved UT3 cleft interacts with ubiquitin C-terminal β-strand. **a,b,** The AlphaFold3 model of UT3 in complex with two ubiquitins (UT3:2Ub) is shown (**a**). UT3, the ridge-bound ubiquitin (Ub^Ridge^), and the cleft-bound ubiquitin (Ub^Cleft^) are depicted in blue, pink, and red, respectively. N and C termini of each polypeptide are indicated. Ub^Ridge^ K48 and Ub^Cleft^ G76 are in vicinity as shown in the close-up view (**b**). **c,** Ufd1 carrying photo-reactive probe Bpa and an HA tag was preincubated with polyubiquitinated sfGFP [Ub(n)-sfGFP], Npl4, ATPase-dead mutant of Cdc48 (E588Q), and ATP, followed by UV-induced photo-crosslinking. The resulting protein samples were analyzed by anti-HA blotting (top) and silver staining (bottom). Asterisk indicates a non-specific band revealed by the anti-HA antibody. **d,** Five ubiquitin β strands are highlighted in colors (PDB: 1UBQ). **e,** GST fusions containing full-length (FL) ubiquitin or indicated ubiquitin fragments were applied in pulldown with HA-tagged Ufd1. Bead-associated proteins were analyzed by anti-HA blotting (top) and Coomassie-blue staining (bottom). Intensities of Ufd1 bands were normalized to input. NS peptide, a non-specific peptide encoded in the pGEX empty vector. **f,** A 2.1-angstrom crystal structure of *Chaetomium thermophilum* UT3 (*ct*UT3; grey) fused to ubiquitin C19 peptide (magenta). **g,** A close-up view of the crystal structure in (**f**). The ubiquitin segment is sandwiched by α4 and the extended β12’ of *ct*UT3. E148 and D194 of *ct*UT3 form hydrogen bonds (orange dashed lines) with ubiquitin residues. All residues are numbered with respect to the corresponding wild-type proteins. **h,** GST pulldown assay was conducted as in (**e**), except with indicated mutant ubiquitin peptides. **i,** Ubiquitin C19 or N20 peptides were immobilized and applied in the pulldown with indicated Ufd1 variants. Bead-associated proteins were analyzed by Coomassie blue staining.

To experimentally confirm the occupancy of UT3 cleft by ubiquitin, we substituted Lys141 or Pro198 of Ufd1, residues at the cleft site, with the Bpa probe (**Fig. 2a**). The Bpa-incorporated Ufd1 proteins were incubated with polyubiquitinated sfGFP, Npl4, and Cdc48, and then irradiated by ultra-violet (UV) to induce Bpa-mediated crosslinking. A significant portion of Ufd1 proteins became slow migrating, smear-like species (**Fig. 2c**; lanes 5 and 6), indicative of covalent crosslinks between UT3 and polyubiquitinated substrates.

The UT3:2Ub model predicts that UT3 cleft associates with a ubiquitin β strand. We then examined interactions between Ufd1 and all five β strands of ubiquitin (**Fig. 2d**). Interestingly, only the glutathione-S-transferase (GST) fused with ubiquitin β5 showed significant Ufd1 binding that is comparable to full-length ubiquitin (**Fig. 2e**). Furthermore, four ubiquitin fragment peptides – N20 (residues 1-20), M1 (residues 21-40), M2 (residues 39-58), and C19 (residues 58-76) – were synthesized and tested for Ufd1 binding (**Extended Data Fig. 3a**). Again, only the β5-containing C19 peptide showed strong binding with Ufd1. This binding was abolished if the amino acid sequence C19 peptide was scrambled (C19scr). Altogether, these results demonstrated that UT3 cleft specifically interacts with ubiquitin β5.

### Structure of UT3 bound with ubiquitin C-terminal peptide

To validate the complex structure of UT3 cleft and ubiquitin β5, a chimera protein with ubiquitin C19 peptide fused to UT3 was generated (UT3-C19). In contrast to UT3, this UT3-C19 fusion failed to interact with resin-immobilized C19 peptide (**Extended Data Fig. 3b**; lane 8), suggesting that the fused C19 likely outcompetes free C19 peptide for the same binding site on UT3. For better crystallization results, the UT3 sequence from a thermophilic yeast, *Chaetomium thermophilum*, was used (*ct*UT3, residues 24-204 of *ct*Ufd1). Notably, this thermophilic yeast shares identical ubiquitin C19 sequence with the human and budding yeast counterparts (**Extended Data Fig. 1c**). A crystal of *ct*UT3-C19 was diffracted at the resolution of 2.1 Å (**Table 1**), and an atomic model was built by molecular replacement (**Fig. 2f**). Each asymmetric unit contains two molecules with very similar conformations, with an r.m.s.d of 0.425 Å (**Extended Data Fig. 3c,d**). The high resolution allows unambiguous side-chain assignment of most *ct*UT3 and ubiquitin residues, especially at the cleft site (**Extended Data Fig. 3e**).

**Table 1.** Data collection and refinement statistics for crystal structure.

|  | ***ct*UT3-C19 (PDB: <u>22ID</u>)** |
| --- | --- |
| **Data collection** |  |
| Space group | C 2 2 21 |
| Cell dimensions |  |
| *a*, *b*, *c* (Å) | 64.35, 67.47, 177.67 |
| $\alpha$, $\beta$, $\gamma$ (°) | 90, 90, 90 |
| Resolution (Å) | 44.42 – 2.14 (2.19 – 2.14) |
| *R*<sub>merge</sub> | 0.146 |
| *I*/σ*I* | 3.18 |
| Completeness (%) | 97.0 |
| Redundancy | 12.3 |
| **Refinement** |  |
| Resolution (Å) | 44.42 – 2.14 |
| No. reflections | 21088 |
| *R*<sub>work</sub> / *R*<sub>free</sub> | 0.246 / 0.288 |
| **Ramachandran plot** |  |
| Outliers | 0.00 % |
| Allowed | 2.86 % |
| Favored | 97.14 % |
| No. atoms | 3325 |
| Protein | 3138 |
| Ligand/ion | 0 |
| Water | 187 |
| *B*-factors | 30.31 |
| Protein | 97.44 |
| Ligand/ion | 120.87 |
| Water | 80.87 |
| R.m.s. deviations |  |
| Bond lengths (Å) | 0.004 |
| Bond angles (°) | 0.711 |

Data were collected from a single crystal. Highest-resolution shell is in parentheses.

An *apo* structure of *ct*UT3 predicted by AlphaFold3 shares almost identical conformation with *ct*UT3 in the crystal structure (**Extended Data Fig. 3f**), except two changes at the cleft. First, ubiquitin binding at the cleft pushes *ct*UT3 α4 slightly away, making the cleft slightly larger. Second, *ct*UT3 β12 becomes longer due to pairing with ubiquitin. In the *apo* form, β12 contains the residues of Ser196 to Phe199 (**Extended Data Figs. 3f and 1e**). In the ubiquitin-bound structures, β12 is extended (referred as β12’ hereafter), starting from either Glu192 or Asp194. The extended β12’ forms an anti-parallel β sheet with ubiquitin C-terminal peptide, resulting in a pairing mechanism called β augmentation. Similar pairing mechanism has also been reported for some HECT (homologous to the E6-AP carboxyl terminus)-type E3 ligases^32^.

Ubiquitin binding to *ct*UT3 involves not only the main-chain hydrogen bonds by β12’ but also the side chains of several key residues on *ct*UT3. For instance, the backbone amine group of ubiquitin Arg72 (hereafter, ubiquitin residues are numbered with respect to their positions in full-length ubiquitin) is coordinated by a hydrogen bond involving a conserved glutamate residue at *ct*UT3 α4 (**Fig. 2g**; Glu148 of *ct*Ufd1, Glu145 of *sc*Ufd1, and Glu143 of *hs*Ufd1). Ubiquitin Arg74 also forms a side-chain hydrogen bond with a conserved aspartate residue on *ct*UT3 β12’ (Asp194 of *ct*Ufd1, Asp191 of *sc*Ufd1, and Asp189 of *hs*Ufd1). In addition, the side chains of ubiquitin Leu71 and Leu73 are wedged into a hydrophobic core in the cleft, which is formed by *ct*UT3 Leu147, Phe151, Met190, Val195, and Val197 (**Extended Data Fig. 3g**). These structural analyses identify the “LRLR” motif at ubiquitin C-terminal tail being critical for its binding with UT3 cleft.

To biochemically verify these interactions, we mutated ubiquitin Leu73 and/or Arg74 and tested their impacts on the cleft binding. Indeed, either L73A or R74E mutant significantly impaired ubiquitin β5 binding to Ufd1 (**Fig. 2h**). To test mutations on the UT3 side, we generated a homology model of *sc*UT3-C19 (**Extended Data Fig. 3h**). Highly conserved residues at the UT3 cleft (**Extended Data Fig. 3i**) were mutated to either alanine or charge-reverse amino acids. Full-length Ufd1 proteins carrying each of these mutations were purified (**Extended Data Fig. 3j**) and subjected to ubiquitin C19 pulldown. Indeed, Ufd1 binding to C19 was abolished by Ufd1 mutations – L144A, E145K or D191K (**Fig. 2i**). In contrast, mutating residues at UT3 ridge, for instance, Leu55 and Glu86, had no effects on Ufd1 binding to C19 (lanes 5 and 8).

### Lys48-linked ubiquitin molecules cooperatively bind to UT3

An AlphaFold model was reported previously for UT3 in complex with one ubiquitin (UT3:1Ub), showing the ubiquitin bound at the ridge site (**Extended Data Fig. 4a**)^21^. Interestingly, the orientations of Ub^Ridge^ in the UT3:1Ub and UT3:2Ub models are different (**Extended Data Fig. 4a,b**), although the conformations of UT3n in the two models appear to be almost identical (**Extended Data Fig. 4c**). For UT3:1Ub, UT3 ridge interacts with ubiquitin β1-β2 hairpin. For UT3:2Ub, however, UT3 ridge engages with ubiquitin Lys48 loop. Thus, UT3 ridge does not have strong selectivity. Nevertheless, the ridge binding in both models appear to involve the Leu55 residue of UT3 and the Ile44 patch of Ub^Ridge^ (**Extended Data Fig. 4a,b**). Consistently, Ufd1 L55A mutant failed to associate with full-length ubiquitin (**Fig. 3a**). Because full-length ubiquitin binding with Ufd1 was diminished only by L55A but not E145K, it implicates that mono-ubiquitin predominantly engages at UT3 ridge, not the cleft.

**Fig. 3:**
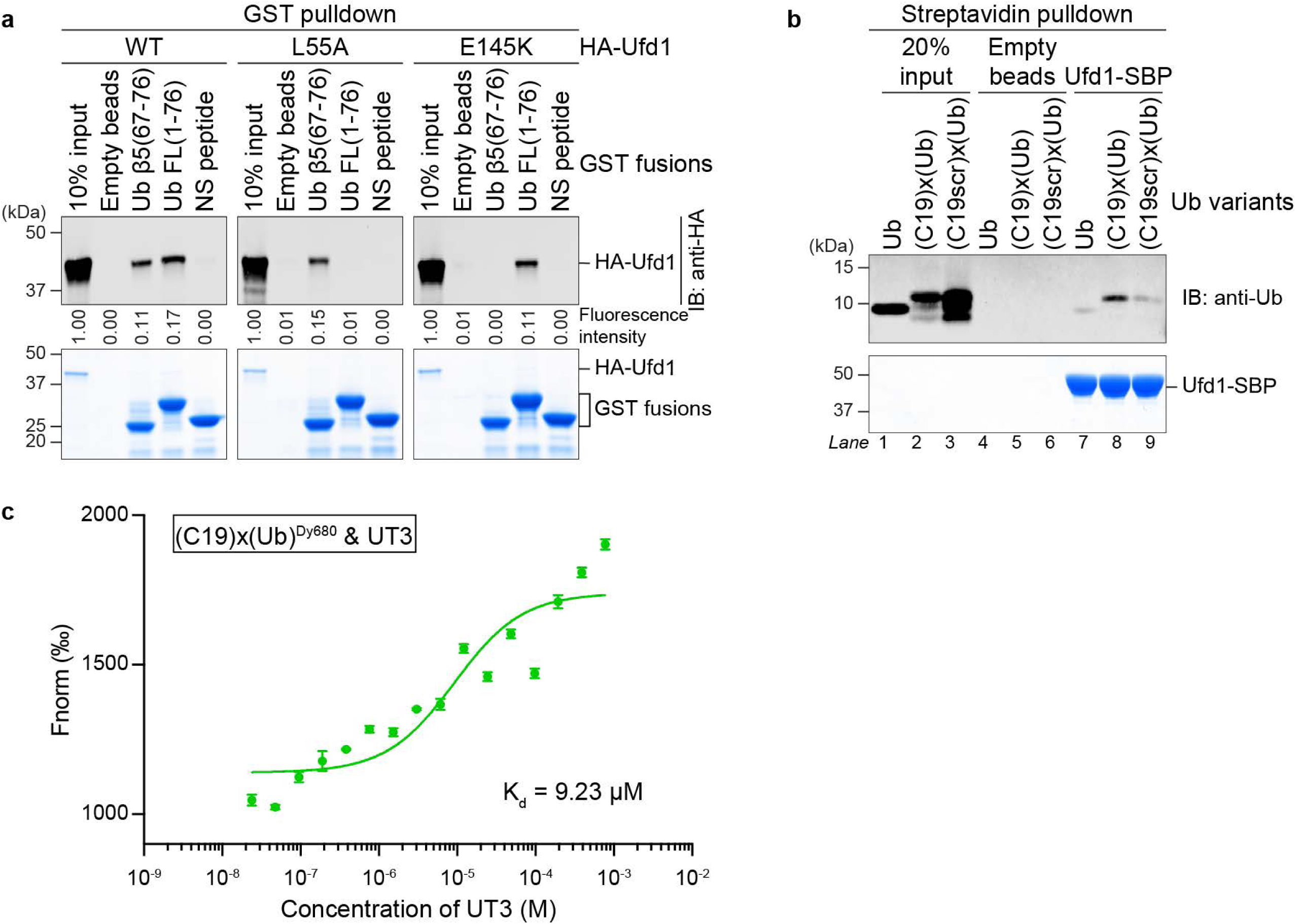
Two K48-linked ubiquitins simultaneously bind to the ridge and cleft sites of UT3. **a,** HA-tagged Ufd1 proteins carrying the ridge mutation L55A or the cleft mutation E145K were applied in GST pulldown by indicated fusion proteins. Bead-associated proteins were analyzed by anti-HA blot (top) and Coomassie blue staining (bottom). Ufd1 band intensities were normalized to input. NS peptide is a non-specific sequence encoded in the pGEX empty vector. **b,** Recombinant ubiquitin or synthetic ubiquitin conjugates carrying C19 peptide [(C19)x(Ub)] or C19-scrambled peptide [(C19scr)x(Ub)] were applied in pulldown by SBP-tagged Ufd1 and streptavidin resin. Bead-associated proteins were analyzed by immunoblotting against ubiquitin (top) and Coomassie blue staining (bottom). **c,** The (C19)x(Ub) conjugate was labeled with the DyLight 680 fluorescent dye (Dy680) and then mixed with different doses of UT3 for affinity measurements using micro-scale thermophoresis. Data from three independent experiments were averaged and fitted to calculate the dissociation constant (K_d_).

Next, we investigated potential cooperativity between the two UT3-bound ubiquitin molecules. Using chemical synthesis, we generated a conjugate of C19 peptide and a full-length ubiquitin (FL Ub), where the C-terminal glycine residue of C19 is linked to the Lys48 residue of the FL Ub via an isopeptide bond [(C19)x(Ub); **Extended Data Fig. 4d-f**]. In this (C19)x(Ub) conjugate, C19 should stay in an unfolded state and is ready to bind with UT3 cleft, while the FL Ub is expected to be well folded and can interact with UT3 ridge. In a pulldown assay by Ufd1, this (C19)x(Ub) conjugate showed stronger binding than ubiquitin alone (**Fig. 3b**). A similar conjugate containing the C19scr peptide and FL Ub [(C19scr)x(Ub)], however, interacted with Ufd1 in the same efficiency as ubiquitin alone. Micro-scale thermophoresis (MST) measured the binding affinity between (C19)x(Ub) and UT3 being 9.23 µM (**Fig. 3c**). In contrast, the affinities of UT3 binding to the (C19scr)x(Ub) conjugate and to the C19 peptide were measured as 73.02 µM and 46.38 µM, respectively (**Extended Data Fig. 4g,h**). Collectively, these lines of evidence implies that Lys48-linked ubiquitin molecules allow simultaneous and cooperative binding to both the ridge and cleft sites of UT3.

### Ubiquitin binding to UT3 cleft triggers its unfolding

We next examined the importance of UT3 ridge and cleft in triggering ubiquitin unfolding. Using the dye accessibility assay, we found that UT3-induced ubiquitin unfolding was largely diminished if either the ridge or the cleft was mutated or blocked (L55A, E145K, or UT3-C19 fusion; **Fig. 4a** and **Extended Data Fig. 5a-c**). Several pieces of evidence suggest that UT3 ridge binding *per se* cannot induce ubiquitin unfolding. First, according to the UT3:1Ub and UT3:2Ub models, ubiquitin remains fully folded when it binds at the ridge site. Second, mono-ubiquitin predominantly binds to UT3 ridge (**Fig. 3a**), but I3C mono-ubiquitin was barely unfolded by UT3 (**Fig. 1d**).

**Fig. 4:**
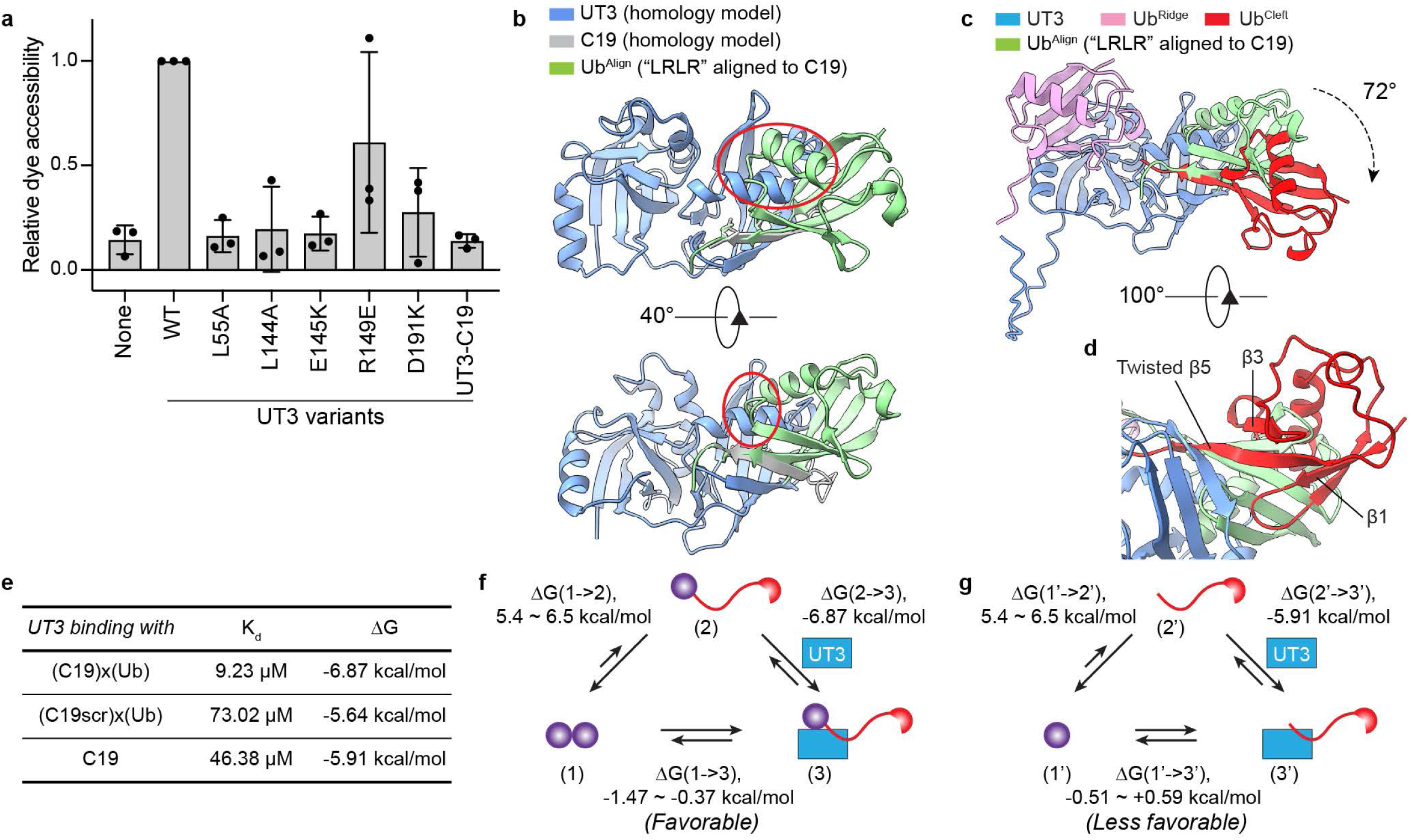
Ubiquitin binding to UT3 cleft induces its unfolding. **a,** Indicated UT3 variants were tested in dye accessibility assay with short-chain I3C polyubiquitins. Intensities of dye fluorescence on I3C di- or tri-ubiquitin were normalized to the WT UT3 samples. The resulting relative dye accessibility scores in three biological replicates were plotted. Error bars show means and s.d. **b,** A folded ubiquitin (green; PDB: 1UBQ) is superimposed to the homology model of UT3-C19 (blue and grey) with the “LRLR” motifs aligned. Shown are two views indicating steric clashes between the aligned, folded ubiquitin (Ub^Align^) and UT3c subdomain (red circles). **c,d,** The UT3:2Ub model is superimposed to the UT3-C19 and Ub^Align^ model in (**b**) with the UT3 domains aligned. Shown in (**c**) are UT3:2Ub (blue, pink, and red) and Ub^Align^ (green). The steric clash between Ub^Align^ and UT3 can be avoided if the residues 1-70 of Ub^Align^ rotates for 72° (dash line). However, the rotation of this ubiquitin leads to a conformationally twisted β5, which may result in the dissociation of β5 from pairing with β1 and β3 (**d**). **e,** A summary of K_d_ and calculated ΔG for UT3 binding to (C19)x(Ub), (C19scr)x(Ub), and the C19 peptide. **f,** A schematic diagram shows three states of di-ubiquitin – fully folded (1), one ubiquitin being unfolded (2), bound to UT3 while one ubiquitin being unfolded (3). Energy for UT3-induced ubiquitin unfolding (1->3) can be calculated from the ΔG for ubiquitin unfolding (1->2) and that for UT3 binding to K48-linked ubiquitin conjugates (2->3). **g,** A schematic diagram similar to (**f**), except with mono-ubiquitin. The three states with mono-ubiquitin are designated as 1’, 2’, and 3’ to distinguish with the di-ubiquitin model in (**f**).

Given the cooperativity between the ridge and cleft in binding to Lys48-linked di-ubiquitin, UT3 ridge probably contributes to ubiquitin unfolding by strengthening ubiquitin interaction with the cleft,.

To understand how the cleft binding leads to ubiquitin unfolding, we superimposed a folded ubiquitin (PDB: 1UBQ) to the UT3-C19 homology model, with the two “LRLR” motifs aligned together. It results in steric clash between the aligned ubiquitin (Ub^Align^) and the UT3c subdomain (**Fig. 4b**). To avoid the clash, Ub^Align^ would have to rotate its residues 1-70 for 72°, while keep its “LRLR” motif bound in the cleft (**Fig. 4c** and **Supplemental Video 1**). Such a rotation allows Ub^Align^ to adopt a conformation similar to Ub^Cleft^ in the UT3:2Ub model. The rotation also causes substantial twist of the β5 strand of Ub^Align^, making the ubiquitin molecule sterically less stable. We propose that Ub^Cleft^ in the UT3:2Ub model represents an energetically unstable, intermediate state. Following the β5 twist, a possible outcome is the break of β5 from pairing with β1 and/or β3 (**Fig. 4d**), resulting in disruption of the β-grasp fold and ubiquitin unfolding. This is consistent with earlier simulation work studying ubiquitin unfolding intermediates, which has found that the disruption of β1-β5 or β5-β3 pairing is the first event during ubiquitin unfolding^33–35^. Taken together, our model suggests that Ufd1 leverages the cleft binding with ubiquitin β5 to cause conformational alteraction and subsequent unfolding of the ubiquitin molecule.

We also investigated whether ubiquitin unfolding is thermodynamically favorable upon UT3 binding. Ubiquitin is perceived to be thermodynamically stable. Earlier experiments concerning the process of ubiquitin unfolding by guanidine hydrochloride have measured the unfolding energy ranging from 5.4 to 6.5 kcal/mol^36–38^. Because the binding affinity between (C19)x(Ub) and UT3 was determined as 9.23 µM, the binding energy ΔG is calculated as -6.87 kcal/mol (**Fig. 4e**). Therefore, the net ΔG for UT3-induced ubiquitin unfolding should range from -1.47 to -0.37 kcal/mol, making the reaction thermodynamically favorable (**Fig. 4f**).

The binding affinity between C19 and UT3 was measured as 46.38 µM, i.e. a ΔG of - 5.91 kcal/mol. If UT3 binds the C-terminal β5 of a mono-ubiquitin at the cleft, the binding energy might not be large enough to compensate for the ubiquitin unfolding energy (**Fig. 4g**). This explains why mono-ubiquitin barely gets unfolded by UT3 (**Fig. 2a**).

### UT3 cleft and ridge sites are important for the function of the Cdc48/p97 ATPase complex

Previous studies have demonstrated that the UT3 domain of Ufd1 is critical for the function of the Cdc48 ATPase complex^17,19^. Here we further examined the importance of UT3 cleft and ridge sites in the assembly of the initiation complex. Point mutations at either UT3 ridge or cleft did not affect Ufd1 binding to Npl4 and Cdc48. But these mutations compromised the recruitment of polyubiquitinated substrates, to various degrees (**Fig. 5a**). Among them, Ufd1 L55A and E145K mutants showed the strongest defects (lanes 3 and 6). Ufd1 T194A and F196A, two mutations at the lower part of UT3 cleft (**Extended Data Fig. 3i**) showed no defects, suggesting the lower cleft is not involved.

**Fig. 5:**
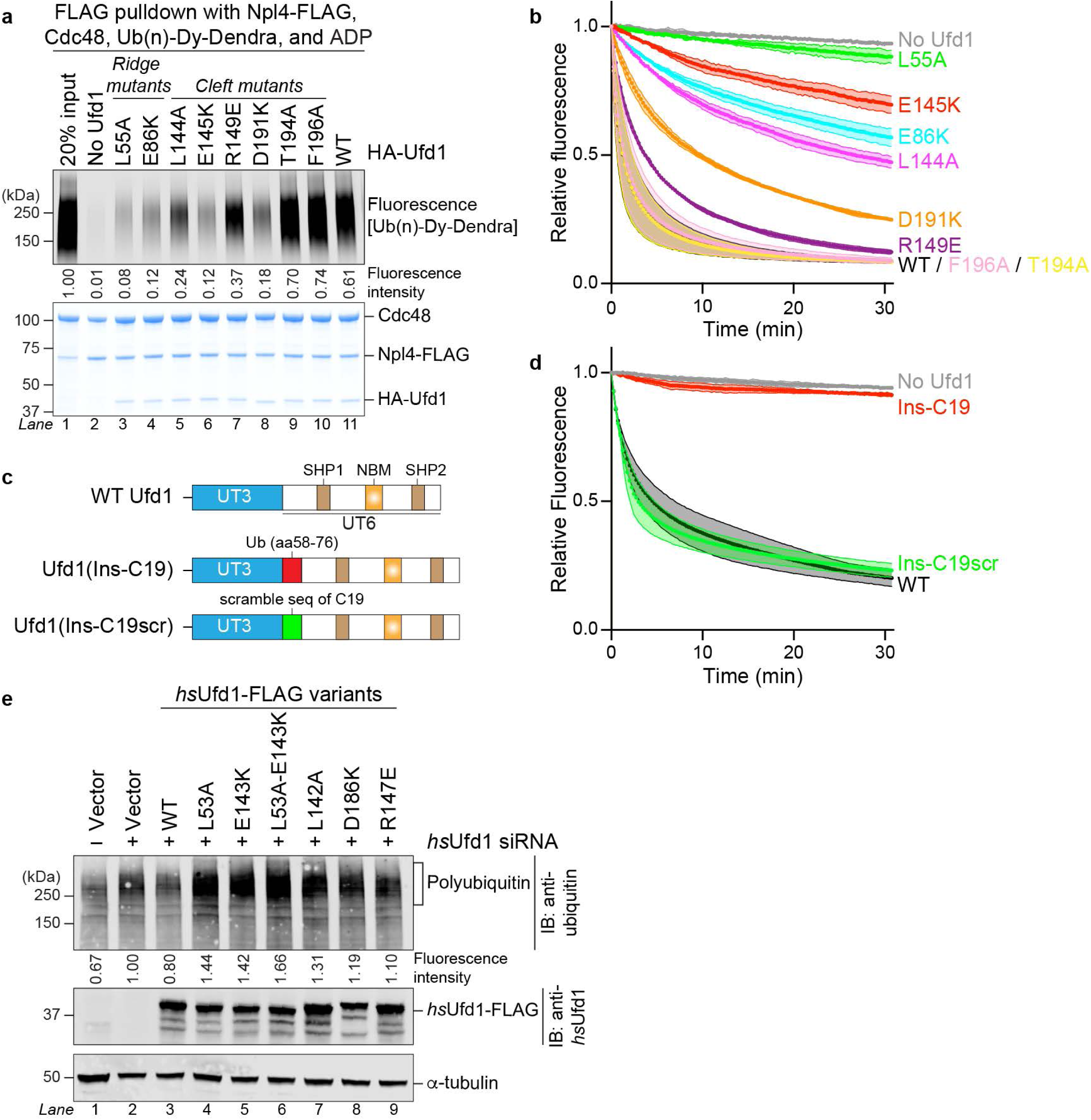
Both UT3 ridge and cleft sites are essential for the functions of Cdc48/p97 ATPase *in vitro* and in cells. **a,** Indicated Ufd1 variants were mixed with polyubiquitinated, dye-labeled Dendra substrate [Ub(n)-Dy-Dendra], Npl4, and Cdc48, followed by anti-FLAG pulldown. Beads-bound proteins were analyzed by dye fluorescence scanning and Coomassie blue staining. Fluorescence intensities were normalized to input. **b,** Indicated Ufd1 variants were tested in Dendra substrate unfolding assay. Intensities of photo-converted Dendra fluorescence were normalized to the zero-timepoint values of each reaction. Data are shown as means +/- s.d. from three independent experiments. Note that the data of WT and T194A samples overlap. **c,d,** C19 or C19-scrambled peptide was inserted between the UT3 and UT6 domains of Ufd1. The resulting chimeras were purified and tested in Dendra substrate unfolding assay in three independent replicates as in (**b**). **e,** Endogenous *hs*Ufd1 in HeLa Tet-On cells were knocked down using specific siRNA, followed by expression with indicated siRNA-resistant variants. Harvested cell lysates were analyzed by immunoblotting against ubiquitin (top), *hs*Ufd1 (middle), and α-tubulin (bottom). Intensities of accumulated polyubiquitinated species were quantified and normalized to the sample in lane 2.

Next, we interrogated the importance of UT3 ridge and cleft in substrate unfolding by the ATPase complex. In the Dendra substrate unfolding assay, Ufd1 L55A and E145K mutants exhibited the strongest defects, whereas T194A and F196A again had no effects (**Fig. 5b**). To rule out the possibility that Ufd1 cleft residues, including E145, are critical for UT3 structure rather than ubiquitin binding, we also employed ubiquitin C19 peptide to block UT3 cleft. Addition of excessive amount of C19, however, only resulted in marginal defects in Dendra unfolding (**Extended Data Fig. 6a**). We reasoned that the mild effect was due to a low occupancy of UT3 cleft by free C19 peptide. To improve the occupancy, C19 peptide was inserted between the UT3 and UT6 domains of Ufd1 (Ins-C19; **Fig. 5c**). Insertion of a TEV cleavage site at the same junction did not affect Ufd1 function^17^. This Ins-C19 variant contains an N-terminal UT3-C19 fusion that is homologous to the crystal structure construct *ct*UT3-C19 (**Fig. 2f**) and is expected to fully block the UT3 cleft site. Indeed, Ufd1(Ins-C19) completely abrogated Dendra unfolding (**Fig. 5d**). In contrast, insertion of the scrambled C19 peptide (Ins-C19scr) did not affect Dendra unfolding at all.

Moreover, we tested if the UT3 ridge and cleft sites are important for human Ufd1 in cells. Consistent with earlier reports^39,40^, knockdown of endogenous human Ufd1 (*hs*Ufd1) using siRNA led to accumulation of high-molecular-weight polyubiquitin species (**Fig. 5e**). Expression of siRNA-resistant wild-type *hs*Ufd1 reversed the accumulation, but expression of the unfolding-defective mutants of Ufd1 failed to rescue. Collectively, these *in-vitro* and *in-vivo* data demonstrate that both Ufd1 ridge and cleft are critical for the Cdc48/p97 ATPase complex in recruiting and unfolding polyubiquitinated substrates, as well as in promoting substrate degradation by the proteasome.

### Full-length Ufd1 facilitates the capture of the unfolded ubiquitin by Npl4 and Cdc48

Our work illustrates that the UT3 domain is sufficient in triggering ubiquitin unfolding. It raises a question about the function of UT6. Earlier studies have demonstrated that UT6 is responsible for binding with Npl4 and Cdc48/p97^24,41^. The importance of UT6 is highlighted by the initiation complex assembly experiment, in which removing UT6 from Ufd1 failed to recruit polyubiquitinated substrates to the Cdc48 ATPase complex (**Extended Data Fig. 2d**). In fact, UT3 itself did not interact with Cdc48 or Npl4.

Moreover, in Dendra substrate unfolding assay, substitution of full-length Ufd1 with split UT3 and UT6 domains abolished substrate unfolding (**Fig. 6a,b**; “equal molar”). This further implies that UT3 tethering to UT6 is important.

**Fig. 6:**
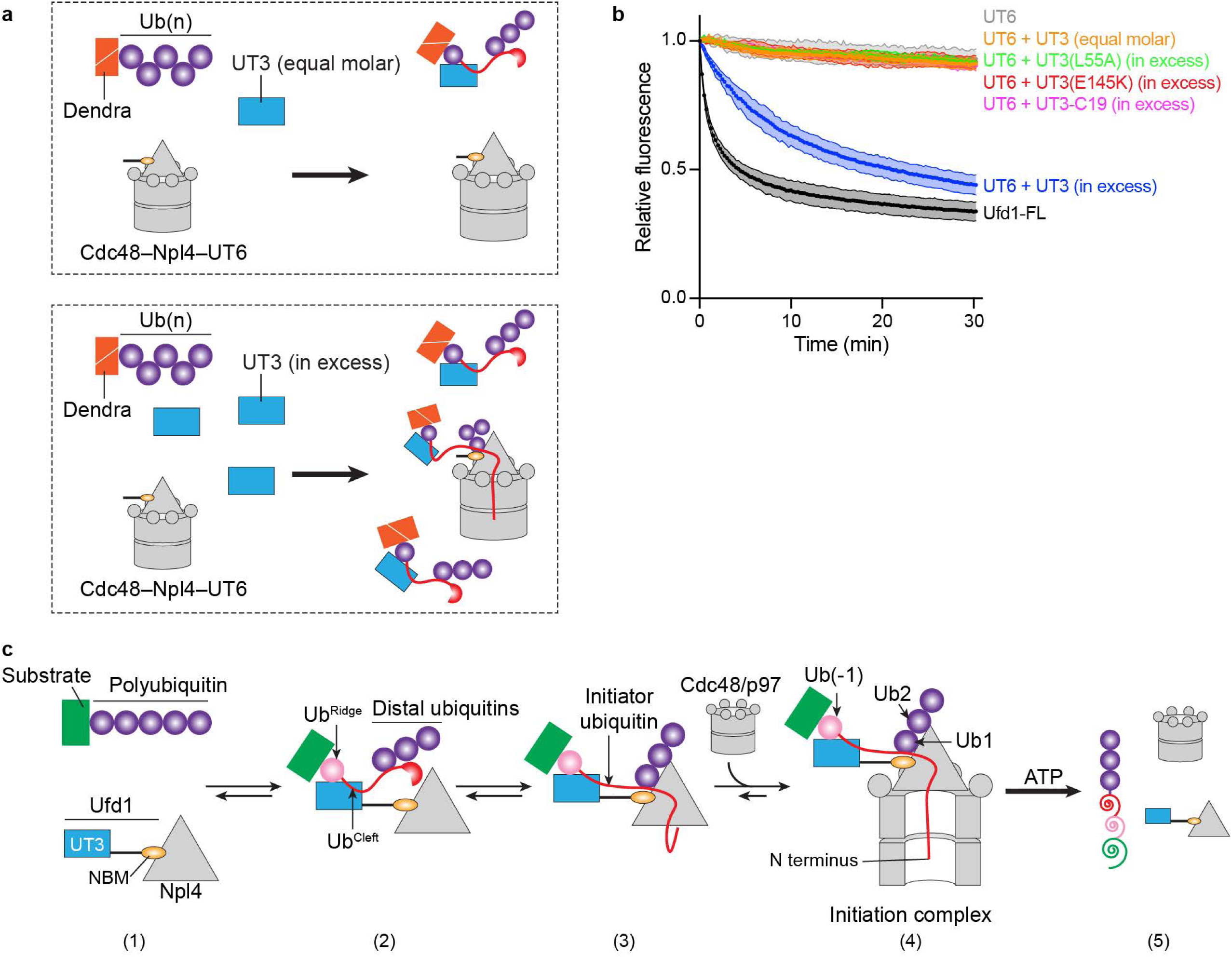
Full-length Ufd1 facilitates the capture of UT3-unfolded ubiquitin by Npl4 and the Cdc48/p97 ATPase. **a,b,** Dendra substrate unfolding assay was performed with split UT3 and UT6 domains. UT3 and indicated variants were added either at equal molar ratio of UT6 or in excess. Data are shown in (**b**) as means +/- s.d. from three independent experiments. **c,** A schematic model delineates the process of ubiquitin unfolded by the UT3 domain of Ufd1 and subsequent ubiquitin capture by Npl4 and Cdc48/p97. Two K48-linked ubiquitin molecules within a polyubiquitin chain is bound by UT3. Ub^Ridge^ remains folded, whereas Ub^Cleft^ gets unfolded. Next, distal ubiquitins interact with Npl4, bringing the unfolded Ub^Cleft^ close to Npl4. Binding of Npl4 to Cdc48/p97 further recruits the unfolded Ub^Cleft^ to the ATPase central pore, leading to the assembly of an initiation complex. Notably, all above-described steps are independent of ATP hydrolysis. Upon ATP hydrolysis, the polyubiquitinated substrate will be unfolded by the ATPase and eventually released.

Then, we tested if UT6 can be replaced by merely an Npl4-binding motif (NBM). Indeed, truncated Ufd1 containing only UT3 and NBM (UT3-NBM) was able to support Dendra unfolding, although at a slightly lower efficiency (**Extended Data Fig. 6b**). In addition, we found that UT3 fused directly to Npl4 (Npl4-UT3), although lacking the entire UT6, was also functional to some extent. These findings are consistent with the hypothesis that UT6 is only important on anchoring UT3 to the ATPase complex.

Surprisingly, substantial Dendra unfolding was observed when the untethered UT3 was added in large excess (250 folds) (**Fig. 6a,b**; “UT3 (in excess)”). This is because excessive UT3 generates more unfolded ubiquitins so that polyubiquitinated substrates are more likely to be captured by the ATPase complex. Consistently, unfolding-defective UT3 variants, UT3 L55R, E145K, or the UT3-C19 fusion, failed to promote Dendra unfolding even if they were added in large excess (**Fig. 6b**). Collectively, these results suggest that the capture of UT3-unfolded ubiquitin by Npl4 and Cdc48 is a rate-limiting step. This capturing step is normally facilitated by anchoring UT3 to the ATPase complex via UT6. In the absence of UT6, ubiquitin capture can be partially restored by tethering UT3 to Npl4 directly or indirectly via an NBM. Alternatively, the function of UT6 can be bypassed by simply adding excessive amount of UT3, which generates more unfolded ubiquitin molecules. Therefore, a key function of the full-length Ufd1 is coupling the generation of unfolded ubiquitin with the ubiquitin capture by Npl4 and Cdc48.

## DISCUSSION

For decades, the function of Ufd1 has been thought to be simply bridging the interaction between polyubiquitinated substrate and the Cdc48/p97 ATPase. Here we uncovered a unique and more fundamental function of Ufd1, which is to induce the unfolding of a ubiquitin molecule within polyubiquitin chains. Our results lead to a model that explains how a ubiquitin molecule is unfolded in an ATP-independent manner and subsequently captured by the Cdc48/p97–Ufd1–Npl4 ATPase complex.

The hexameric Cdc48/p97 ATPase forms stable complex with the heterodimer of Npl4 and Ufd1. This ATPase complex can recruit and process substrates carrying a Lys48-linked polyubiquitin chain. First, two adjacent ubiquitin molecules within the polyubiquitin chain bind simultaneously to the UT3 domain of Ufd1. The proximal molecule at the Lys48 side of the isopeptide bond binds at UT3 ridge, while the distal molecule at the Gly76 side of the isopeptide bond docks at UT3 cleft via its C-terminal “LRLR” motif. Key UT3 elements for the cleft binding include hydrophobic residues at the cleft, the extended β12’, and a conserved glutamate residue at UT3 α4. Residues 1-70 of the cleft-bound ubiquitin (Ub^Cleft^) would undergo a 72° rotation to avoid steric clash, leading to a conformational distortion of the β5 strand of Ub^Cleft^ (**Supplemental Video 1**). Such a distorted configuration is not stable, causing disengagement of β5 from pairing with β1 and/or β3 that eventually results in the unfolding of the Ub^Cleft^ molecule (**Fig. 6c**, step 2).

Next, ubiquitin molecules distal to Ub^Cleft^ interacts with Npl4 and brings the middle segment of Ub^Cleft^ in vicinity to the Npl4 groove (**Fig. 6c**, step 3). Binding of the unfolded ubiquitin to Npl4 further projects the ubiquitin N-terminal segment to the central pore of Cdc48/p97 (**Fig. 6c**, step 4). As a result, one ubiquitin molecule is unfolded by UT3 and further stabilized by Npl4 and Cdc48/p97, leading to the assembly of the initiation complex. Notably, this process of ubiquitin unfolding does not requires ATP hydrolysis.

A standing question regarding the initation complex is which molecule(s) along a polyubiquitin chain can serve as the initiator ubiquitin. Previous FRET experiments revealed that fluorophore attached to the first ubiquitin at the promixal end (i.e. the ubiquitin directly attached to the substrate) could not be the initiator^21^. Our current model provides a perfect explanation to this observation. When a ubiquitin molecule binds to UT3 cleft and gets unfolded, it requires at least one more ubiquitin at its proximal side to engage with the UT3 ridge. The first ubiquitin at the proximal end of the chain cannot bind to UT3 cleft, thus cannot serve as the initiator. In addition, if the last two ubiquitins at the distal end of a polyubiquitin chain are unfolded by UT3, they would not have enough distal ubiquitin molecules to engage with the Npl4 tower^15^. As a consequence, the unfolded ubiquitin could not engage with Npl4 or Cdc48, and would likely refold. Other than these terminal molecules, any ubiquitin in between can presumably be unfolded by UT3 and captured by the ATPase complex as the initiator, as suggested by previous HDX/MS results^17^.

It is known that the Cdc48/p97 ATPase complex only recruits substrates carrying a Lys48-linked polyubiquitin chain. The linkage specificity is deciphered at the first step, as UT3 only acts on Lys48-linked di-ubiquitin (**Extended Data Fig. 2a**). Later, this selectivity is further reinforced by polyubiquitin binding to Npl4^15^. As a result, the Cdc48/p97 ATPase complex exhibits strong specificity towards Lys48-linked polyubiquitin chains.

Although ubiquitin is remarkably stable, earlier measurements and our analysis found that the energy of UT3 ridge and cleft binding to two ubiquitin molecules is enough to compensate for ubiquitin unfolding energy. Furthermore, ubiquitin molecules in a polyubiquitin chain have lower unfolding energy than mono-ubiquitin, making the UT3-induced ubiquitin unfolding more favorable (**Fig. 4f**). This is because attachment of even a single ubiquitin is sufficient to destabilize the modified protein^42,43^. Except the molecule at the distal end, all ubiquitins within a polyubiquitin chain are modified by another ubiquitin. In addition, melting temperature measurements have revealed that as polyubiquitin chain grows longer, the ubiquitin molecules become less stable^44^.

Therefore, long-chain polyubiquitins have higher probability of being unfolded by UT3 and being the substrate for the ATPase complex. Consistent to this model, polyubiquitin species accumulated by p97 inhibition migrated more slowly than those accumulated upon proteasome inhibition^10^.

Di-ubiquitin binding to UT3 is not very strong (**Fig. 3c**), indicating that they may dissociate after ubiquitin unfolding. Following the dissociation, unfolded Ub^Cleft^ could refold, if it is not already captured by Npl4 or Cdc48/p97. This could be important for cell to avoid accumulation of unfolded ubiquitin molecules. Protein aggregates in neurodegenerative diseases often contain ubiquitin^45,46^. Further analyses are needed to investigate if these ubiquitin molecules are in the unfolded form, and if the Ufd1 function is connected to the ubiquitin accumulation in protein aggregates.

Our results suggest that the key roles of the UT6 domain and Npl4 are to bring together the UT3-unfolded ubiquitin and Cdc48/p97 ATPase. It is worth noting that UT3 adopts a strikingly similar structure as the N domain of Cdc48/p97. However, full-length Cdc48 containing the N domain cannot unfold I3C di-ubiquitin (**Fig. 1c**, lane 5). In fact, Cdc48/p97 N domain employs its cleft site for binding with the SHP motifs within Ufd1 UT6^41^. A plausible speculation is that in early evolution, Cdc48/p97 ancestor uses its own N domain to capture and unfold ubiquitins; the unfolded ubiquitin is then projected into the ATPase central pore for processing. During evolution, the ubiquitin unfolding function is inherited by Ufd1, a putative gene duplicate from the primordial Cdc48/p97; Ufd1 remains anchored to the ATPase through binding to the ATPase N domain. Meanwhile, the ATPase largely expands its binding partners via the N domain and participates in various cellular functions beyond polyubiquitin processing.

Our structural and biochemical analyses imply that ubiquitin C-terminal “LRLR” motif plays a key role in UT3 cleft binding. Interestingly, two other ubiquitin-like proteins, Nedd8 and ISG15, also contain a “LRLR” or similar motif at their C-termini (**Extended Data Fig. 3k**). Although no evidence shows these two ubiquitin-like proteins can be linked into polymers, it remains unclear if they can interact with UT3 on their own, or with a modified substrate occupying UT3 ridge. Further investigations are needed to explore any functional links of Ufd1 to Nedd8 and/or ISG15.

## ACKNOWLEDGMENTS

We thank the staff of the BL-10U2 beamline at the National Center for Protein Sciences Shanghai at Shanghai Synchrotron Radiation Facility for support in X-ray diffraction and data collection, Yifei Zhao and Jiao Liu at the Protein Characterization and Crystallography Facility of Westlake University for assistance in crystallization and data processing, the High-throughput Screen Facility at Westlake University for use of the plate-reader equipment. We are grateful to Lei Liu for advice on the synthesis of ubiquitin conjugates, and to Shutao Qi and Hongtao Yu for sharing the HeLa TetOn cell line. We thank Tom A. Rapoport and Hongtao Yu for critical reading on the manuscript. This work was supported by the Key R&D Program of Zhejiang (grant no. 2024SSYS0029).

## AUTHOR CONTRIBUTIONS

Z.J. conceived and supervised the project. Y.W. performed most of the biochemical and biophysical experiments. Z.Z. conducted the human cell complementation analysis. W.H. synthesized the ubiquitin conjugates and conducted the QC validation analysis. P.W. helped with the data collection for crystal diffraction. Z.J. wrote the manuscript with the inputs from all the others.

## DECLARATION OF INTERESTS

The authors declare no competing interests.

### Data and Code Availability

Coordinates and structure files have been deposited in the Protein Data Bank (PDB) under accession number 22ID (ctUT3-C19). This paper does not report original code. Any additional information required to reanalyze the data reported in this paper is available from the lead contact upon request.

## METHODS

### Bacteria cultures

Bacterial expressing plasmids were transformed into *Escherichia coli* BL21 CodonPlus (DE3) RIPL cells (Agilent), unless stated otherwise. Selected clones were grown in Terrific Broth (BD) to an OD_600_ of 0.8. Protein expression was induced by addition of 0.1 mM isopropyl b-D-1-thiogalactopyranoside (IPTG). Bacterial cultures were continued at 16°C for 16 hours before harvest.

For expression of the Bpa-incorporated proteins, the expression plasmids were transformed into *E. coli* BL21 (DE3) TSR2566 Chemically Competent Cell (Tsingke) harboring the plasmid pEVOL-pBpF^47^. Selected clones were grown in Terrific Broth to an OD_600_ of 0.8. Protein expression was induced by the addition of 0.2% L-arabinose, 0.5 mM Bpa, and 0.1 mM IPTG, and the incubation was continued at 16°C for 16 hours before harvest.

### Yeast cultures

Plasmids encoding *Saccharomyces cerevisiae* Ubr1 were transformed into the INVSc1 yeast strain (Thermo). Selected yeast clone was grown in synthetic dropout (SD) medium for 24 hours, and then supplemented with yeast culturing medium containing 2% galactose to induce protein expression. After 24 hours, the cell cultures were harvested.

### Cell cultures and transfection

HeLa Tet-On 3G cells (Clontech, cat#630901) were cultured in Dulbecco’s Modified Eagle Medium (DMEM, Invitrogen) supplemented with 10% fetal bovine serum (Thermo) and 2 mM L-glutamine (Thermo). For RNAi experiments, cells were transfected with siRNA oligonucleotides (Genepharma) with Lipofectamine RNAiMAX (Thermo) and harvested at 24-72 hours after transfection. Sequence of the *hs*Ufd1 siRNA is (5’ to 3’) GCACUUAGGAACUUUGCCU-dTdT.

### Plasmids

For human cell experiments, the wild-type gene of *Homo sapiens* Ufd1 (*hs*Ufd1) was cloned into a modified pcDNA3.1 backbone using the NotI and AscI restriction enzyme sites. Silent mutations were introduced to the siRNA-targeting region, in order to make the transgene resistant to *hs*Ufd1 siRNA. A sequence encoding the 3xFLAG tag (“DYKDHDGDYKDHDIDYKDDDDK”) was inserted at the C-terminus of each gene to express FLAG-tagged *hs*Ufd1. Mutations were introduced by overlapping PCR.

For bacterial expression of full-length (FL) ubiquitin, a pK27(His14-SUMO-*hs*Ub) vector containing the pK27 plasmid backbone, an N-terminal His14-SUMO (small ubiquitin-like modifier) tag, and a human ubiquitin sequence was used. Wild-type *S. cerevisiae* ubiquitin (*sc*Ub) and the I3C mutant were cloned into the pK27(His14-SUMO-*hs*Ub) vector using the Gibson assembly method. Note that treatment of the His14-SUMO-*hs*Ub-*sc*Ub protein with a de-ubiquitinating enzyme (DUB) would cleave off only the yeast ubiquitin, not the human ubiquitin. For M1-linked I3C di-ubiquitin, tandem repeats of I3C ubiquitin was cloned into a pK27(His14-SUMO-Ala) vector via Gibson assembly method. The insertion of an alanine residure between the SUMO tag and ubiquitin allows efficient cleavage of di-ubiquitin using a SUMO protease. For GST pulldown experiments, ubiquitin fragments and FL ubiquitin were cloned into a modified pGEX6p1 vector using NotI and AscI sites. The resulting linker between the GST tag and ubiquitin sequences is “DHPPKSDLEVLFQGPLGSGAAA”, which contains an HRV-3C protease cleavage site. The empty pGEX6p1 vector encodes the GST tag, linker, and a 12-residue, non-specific (NS) peptide, “RPELGAPLEAAS”.

Fluorescent model substrates containing super-folder GFP (sfGFP) or photo-convertible moxDendra2 (Dendra) were constructed as previously described^17^. Briefly, a 43-residue N-end-rule degron (NeD) was fused to the N-terminus of either fluorescent protein. The fusion constructs were then cloned into the pK27(His14-SUMO) vector using Gibson assembly method. The NeD sequence is as follows (note that an N-terminal arginine is generated after SUMO protease cleavage): “RHGSG<u>C</u>GAWLLPVSLVKRKTTLAPNTQTASPPSYRALADSLMQ”. The cysteine residue is used to react with fluorescent dye-conjugated maleimide moieties. The mouse Ube1 gene was cloned into a pET28 vector using NdeI and EcoR1 sites as previously described^48^. The coding region of gp78^RING^-Ube2g2 fusion, encoding the RING domain of human gp78 (residues 322-393) and human Ube2g2 with the linker sequence GTGSH in between, was constructed as described previously ^7^ and cloned into the pET28(His6-TEV) vector using NotI and AscI sites. *S. cerevisiae* Ubc2, Ubc4, and Rsp5 genes were cloned into pK27(His14-SUMO) vector using Gibson assembly method. The catalytic domain of human Usp2 (Usp2^cat^, residue 251-605) carrying two mutations, E472A and K473A was cloned into a pET28 vector using Gibson assembly method.

All Ufd1 variants were cloned into the pK27(His14-SUMO) vector using the NotI and AscI sites. A sequence encoding the hemagglutinin (HA)-tag (“YPYDVPDYA”) was inserted between the SUMO and Ufd1 sequences, where indicated. A streptavidin-binding protein (SBP)-tag sequence was appended after the Ufd1 sequence as appropriate. UT3 contains the first 201 residues of *S. cerevisiae* Ufd1. *ct*UT3 contains residues 24-204 of *C. thermophilum* Ufd1. UT3-C19 and ctUT3-C19 have the ubiquitin C19 (residues 58-76) fused to the C-terminus of the corresponding UT3 domain. UT6 (Ufd1ΔUT3) contains a truncation of the first 200 residues of Ufd1. For UBA/UBL-UT6 chimeras, Dcn1^UBA^ (residues 14-51), Ddi1^UBL^ (residues 1-80), Dsk2^UBA^ (residues 320-373), Ede1^UBA^ (residues 1339-1381), Rad23^UBA1^ (residues 141-189), or Rad23^UBA2^ (residues 356-398) was fused to the N-terminus of UT6. Ufd1(Ins-C19) or Ufd1(Ins-C19scr) have inserted the ubiquitin C19 sequence (residues 58-76) of ubiquitin or a scrambled C19 sequence between the residues 201 and 202 of Ufd1. The amino acid sequence of C19scr is “GLNRLESTGLYKHLIQDVR”. Point mutations, chimera fusions, and insertions were generated by overlapping PCR.

Yeast Npl4 gene was cloned into the pET21 vector using NdeI and AscI sites, with a C-terminal FLAG-His6 tag. Wild-type Cdc48 and its variants were cloned into the pET28 vector using NotI and AscI sites, with a His6-tag and a TEV-protease cleavage site at the N-terminus. Cdc48^E588Q^ has mutated the Walker B motif of the D2 domain and abrogated its D2 ATPase activity. Cdc48^ND1^ contains residues 1-480 of Cdc48. A sequence encoding the FLAG tag was added at the C-terminus of Cdc48, where indicated.

For the expression of p-Benzoyl-phenylalanine (Bpa)-incorporated proteins, residues that are eventually mutated to p-Benzoyl-phenylalanine (Bpa) were mutated to *amber* stop codon (“UAG”) using overlapping PCR. The pEVOL-pBpF plasmid used to produce Bpa-incorporated proteins was a gift from Peter Schultz (Addgene plasmid # 31190).

For yeast expression, *S. cerevisiae* Ubr1 gene was cloned into pRS426Gal1(His14-TEV), using NotI and AscI sites, resulting the N-terminal sequence, “MSKHHHHSGHHHTGHHHHSGSHHHG-ENLYFQ-GAAA”.

### Synthetic ubiquitin peptides

The chemically synthesized ubiquitin N20, M1, M2, and C19 peptides (Synpeptide) contain sequences of *S. cerevisiae* ubiquitin corresponding to residues 1-20, 21-40, 39-58, and 58-76. Sequence of the C19scr peptide is “GLNRLESTGLYKHLIQDVR”. All synthetic ubiquitin peptides also contain a C-terminal cysteine for covalently coupling to SulfoLink resins (Thermo). Fluorescein isothiocyanate (FITC) was added to the N-terminus of synthetic peptide as appropriate.

### Immunoblotting and antibodies

Primary antibodies used in this study are: anti-ubiquitin (Proteintech, cat#10201-2-AP, 1:1000), anti-HA (Cell Signaling Technology, clone C29F4, cat#3724, 1:2000), anti-*hs*Ufd1 (Bethyl, cat#A301-875A, 1:2000), anti-α-tubulin (Cell Signaling Technology, clone DM1A, cat#3873, 1:2000), anti-FLAG (Sigma, clone M2, cat#F1804, 1:2000). Secondary antibodies used in this study are: donkey anti-mouse IgG DyLight 800 conjugated (Thermo Fisher, cat#SA5-10172, 1:5000), donkey anti-mouse IgG DyLight 680 conjugated (Thermo Fisher, cat#SA5-10170, 1:5000), donkey anti-rabbit IgG DyLight 800 conjugated (Thermo Fisher, cat#SA5-10044, 1:5000), donkey anti-rabbit IgG DyLight 680 conjugated (Thermo Fisher, cat#SA5-10042, 1:5000), donkey anti-mouse IgG(H&L) horseradish peroxidase (HRP) conjugated (Thermo Fisher, cat#A16017, 1:5000). The substrate for HRP conjugated secondary antibody was Western Lighting Ultra (Perkin Elmer, NEL111001EA).

### Protein purifications

Cdc48, Npl4, and their variants were expressed and purified as previously described^17^. Briefly, bacterial cells were harvested by centrifugation at 5000 g for 10 min and resuspended in wash buffer (50 mM Tris-HCl, pH 8, 320 mM NaCl, 5 mM MgCl_2_, 10 mM imidazole, 0.5 mM ATP) supplemented with phenylmethylsulfonyl fluoride (PMSF; 1 mM), a protease inhibitor cocktail (APExBio), and DNase I (5 µg/mL). The cells were lysed by sonication or by a high-pressure cell homogenizer (Duoning). Cell lysates were cleared by ultracentrifugation in a Ti-45 rotor (Beckman) at 40,000 rpm for 30 min at 4°C. The supernatants were incubated with Ni-NTA resin (Thermo) that was pre-equilibrated with wash buffer, for 60 min at 4°C. The resin was washed three times with 30 column volumes of wash buffer. Proteins were eluted with elution buffer (50 mM Tris-HCl, pH 8, 150 mM NaCl, 5 mM MgCl_2_, 400 mM imidazole). Relevant fractions were pooled and diluted, before being loaded onto a ReSource Q column (Cytiva) using an AKTApure system (Cytiva). Proteins bound on the ReSource Q column were eluted by a gradient of 50-500 mM NaCl. Relevant fractions were concentrated and then loaded on a gel filtration column, which was equilibrated by size-exclusion chromatography (SEC) buffer (50 mM HEPES, pH 7.4, 150 mM NaCl, 5 mM MgCl_2_, and 0.5 mM tris(2-carboxyethyl)phosphine (TCEP)). Fractions that contain the target proteins were pooled and snap frozen.

Full-length Ufd1, UT3, UT6, and their variants with an N-terminal His14-SUMO tag were purified by Ni-NTA resin as described above. The SUMO protease Ulp1 was added to the eluted Ufd1 proteins and dialyzed against wash buffer containing 10 mM imidazole. The Ulp1-treated samples were incubated with Ni-NTA resin to remove the His14-SUMO tag, and the unbound proteins were concentrated and loaded onto a Superdex 200 Increase column equilibrated with the SEC buffer. Ufd1 variants with incorporated Bpa and *ct*UT3-C19 were purified using the same protocol.

WT ubiquitin and the I3C mutant were purified similar to the Ufd1 protein, except that Usp2^cat^ was used instead of Ulp1 to remove His14-SUMO-*hs*Ub tag and that Superdex 75 Increase column was used for gel filtration. M1-linked I3C di-ubiquitin was expressed and purified as mono-ubiquitin, except that a SUMO protease was used to remove the His14-SUMO tag. GST tagged ubiquitin fragments were purified via GS4B resin (Cytiva). After being eluted by reduced glutathione, the GST-tagged proteins were further purified with a Superdex 200 Increase column equilibrated with the SEC buffer.

Fluorescent substrates (Dendra or sfGFP) containing the N-end rule degron and His14-SUMO tag were purified by Ni-NTA resin, followed by the SUMO tag cleavage, SUMO tag removal with Ni-NTA resin, and gel filtration, similarly to the purification of Ufd1 proteins.

*Mus musculus* Ube1 (mUbe1), the gp78^RING^-Ube2g2 fusion, Ubc2, His14-Ubr1, and Usp2^cat^ were purified by Ni-NTA and size-exclusion chromatography as previously described^17^. Ubc4 and Rsp5 were expressed and purified in the same way as Ufd1 proteins, omitting the His-SUMO tag removal. After being eluted from Ni-NTA resin, the resulting proteins were buffer-exchanged into SEC buffer, concentrated, and snap-frozen.

### Generation of short-chain Ub^I3C^ polymers

60 µM Ub^I3C^ was incubated with 0.8 µM mUbe1, 10 µM gp78^RING^-Ube2g2, and 10 mM ATP for 60 min at 30°C in the ubiquitination buffer (50 mM Tris pH 8.0, 150 mM NaCl, 10 mM MgCl_2_, 2 mM dithiothreitol (DTT)). The protein reaction was then acidified using 0.3 volume of glacier acetic acid. Protein precipitates were removed by centrifugation at 4000 g for 10 min. Next, the supernatant was applied to a ReSource S column (Cytiva). Bound proteins were eluted via a salt gradient of 50-450 mM. Fractions were analyzed by SDS-PAGE. Relevant fractions were buffer exchanged into SEC buffer omitting TCEP.

To generate K63-linked I3C di-ubiquitin, 100 µM Ub^I3C^ was incubated with 800 nM mUbe1, 15 µM Ubc4, 2.5 µM Rsp5, and 5 mM ATP for 60 to 90 min at 30°C in the ubiquitination buffer. The products were then purified via ReSource S column as described above. M1-linked I3C di-ubiquitin was expressed recombinantly.

### Dye accessibility assay

10 µM of the isolated short-chain Ub^I3C^ was incubated with 20 µM UT3, Ufd1, or indicated proteins on ice for 10 min. Then DyLight 800 dye-conjugated maleimide moiety (Thermo) was added and incubated for 2 min. The reactions were quenched by 20 mM dithiothreitol (DTT). The samples were mixed with SDS sample buffer and analyzed by SDS-PAGE, followed by dye fluorescence scanning on an Odyssey imager (LI-COR) and Coomassie blue staining.

### Dye labeling of substrates containing the N-end rule degron

The purified model substrates were reduced with 10 mM TCEP and then incubated with a 3-fold molar excess of DyLight dye-conjugated maleimide moieties (Thermo). The reactions were kept in the dark at room temperature for 2 hours before quenching with 20 mM DTT. The unreacted free dyes were removed by Dye Removal columns (Thermo, cat#22858).

### Photoconversion of substrates containing Dendra

The purified Dendra proteins (4∼8 mg/mL) were placed in a 200-µl PCR tube in an ice bath. A long-wavelength UV flashlight (395-410 nm, DULEX) was positioned 5 cm above the tube, and the sample was irradiated for 1 hour, with occasional mixing.

### Ubiquitination of model substrates

Ubiquitination of the model substrates containing an N-end rule degron was carried out as previously described^17^, with some modifications. Substrate (5 µM) was incubated with *S. cerevisiae* ubiquitin (250 µM), mUbe1 (800 nM), Ubc2 (2 µM), Ubr1 (800 nM), and ATP (10 mM) for 60 min at 30°C in ubiquitination buffer (50 mM Tris pH 8.0, 150 mM NaCl, 10 mM MgCl_2_, 1 mM DTT). The samples were concentrated and loaded onto a Superdex 200 Increase column in SEC buffer. After analysis by SDS-PAGE, fractions containing the desired ubiquitin chain lengths were pooled and snap-frozen. The concentration of the pooled polyubiquitinated substrate was determined with a fluorescence microplate reader, using the non-ubiquitinated substrates as standards. The majority of the final product contained polyubiquitin chains of 10-25 ubiquitin molecules.

### In vitro pulldown experiments

For *in-vitro* pulldown of the initiation complexes, 1 µM polyubiquitinated model substrate was mixed with 1 µM Cdc48, 1 µM Ufd1, and/or 1 µM Npl4 in binding buffer (50 mM Tris-HCl, pH 8, 150 mM NaCl, 10 mM MgCl_2_, and 1 mM DTT) supplemented with 10 mM of ADP. 50 µL of the protein mixture were then incubated with 8 µL pre-equilibrated FLAG antibody M2 agarose beads (Sigma) at 4°C for 1 hour. The beads were then washed three times with binding buffer containing ADP. Bead-bound proteins were eluted with 25 µL of binding buffer supplemented with 0.05% Tween-20 and 0.2 mg/mL 3xFLAG peptide (Synpeptide). The eluted samples were separated on SDS-PAGE, followed by fluorescence scanning on an Odyssey imager (LI-COR) and Coomassie blue staining.

For GST pulldowns with ubiquitin fragments, 30 µg of GST-tagged ubiquitin proteins were mixed with the desired HA-Ufd1 variants in binding buffer (50 mM HEPES-Na, pH 7.4, 150 mM NaCl, 5 mM MgCl_2_, and 0.5 mM TCEP). The beads were then washed three times with binding buffer, and eluted by SDS sample buffer. The eluted proteins were separated on SDS-PAGE, followed by immunoblotting using anti-HA antibody and Coomassie blue staining.

For ubiquitin-peptide pulldown, the synthetic ubiquitin peptides containing a C-terminal cysteine residue were immobilized on the SulfoLink Resin (Thermo) according to the manufacturer’s instructions. Then non-reacted binding sites on the resin were blocked by addition of L-cysteine. Peptide-coupled beads were pre-equilibrated into the protein binding buffer (50 mM HEPES-Na, pH 7.4, 150 mM NaCl, 5 mM MgCl_2_, and 0.5 mM TCEP). Then the beads were incubated with HA-tagged Ufd1, UT3, or indicated variants. Unbound proteins were washed off by the protein binding buffer. Beads-associated proteins were separated on SDS-PAGE and analyzed by Coomassie blue staining and immunoblotting against the HA epitope.

For streptavidin pulldowns, 1 µM of synthetic (C19)x(Ub) or (C19scr)x(Ub) conjugates, or recombinant ubiquitin were incubated with empty streptavidin beads or beads bound with Ufd1-SBP in binding buffer (50 mM HEPES-Na, pH 7.4, 150 mM NaCl, 5 mM MgCl_2_, and 0.5 mM TCEP). The beads were then washed three times with binding buffer, and eluted by SDS sample buffer. The eluted proteins were separated on SDS-PAGE, followed by immunoblotting using anti-ubiquitin antibody and Coomassie blue staining.

### Substrate unfolding assays

With the exceptions mentioned below, substrate unfolding experiments were performed as previously described^17^. Briefly, 400 nM of the polyubiquitinated, photoconverted Dendra proteins were mixed with 400 nM Ufd1 variants, 400 nM Npl4, and 400 nM Cdc48 hexamer in 50 mM HEPES pH 7.5, 150 mM NaCl, 10 mM MgCl_2_, 0.5 mM TCEP, and 0.5 mg/mL protease-free bovine serum albumin (BSA; Sigma). After a 10-min pre-incubation at 30°C, an ATP regeneration mixture was added (10 mM ATP, 20 mM phosphocreatine, 100 µg/mL creatine kinase), and the fluorescence (excitation, 540 nm; emission, 580 nm; gain, 60 to 80) was measured at 15-s intervals for around 30 min, using a Spark Microplate Reader (Tecan) or a Synergy Neo2 Multi-mode reader (BioTek). For each experiment, an empty well was included to determine background fluorescence. The relative fluorescence at time *t* was calculated as [(fluorescence at *t*) – (background fluorescence at *t*)] / [(fluorescence at *t_0_*) – (background fluorescence at *t_0_*)]. Calculated relative fluorescence values were plotted against time using Prism software (GraphPad).

### Photo-crosslinking experiments

The Cdc48-Bpa photo-crosslinking experiments were performed as described^15^, with some modifications. Briefly, Cdc48_ND1^D324Bpa^-FLAG (200 nM), Ufd1 (500 nM), Npl4 (500 nM), and Ub(n)-Dy-sfGFP (1 µM) were mixed in reaction buffer (50 mM HEPES, pH 7.5, 150 mM NaCl, 10 mM MgCl_2_, 0.5 mM TCEP, 0.5 mg/mL protease-free BSA) supplemented with 10 mM of ADP. The reactions were assembled on ice, incubated at 30°C for 10 min, and transferred to individual PCR tubes. A long-wave UV lamp (Blak-Ray) was positioned 5 cm above the tubes, and the samples were irradiated for 30 min. To prevent overheating, an ice-cold metal block was placed in contact with the bottom of the PCR tubes. After irradiation, the samples were diluted 10-fold in dissociation buffer (50 mM Tris-HCl, pH 8.0, 800 mM NaCl, 1% (v/v) Triton X-100, 1 mM EDTA, 0.5 mM DTT) and incubated at room temperature for 5 min. The samples were then applied to 8 µL FLAG antibody M2 magnetic beads (Sigma) equilibrated in dissociation buffer for 1 hour at room temperature. The beads were washed three times, and bound protein was eluted in 50 mM HEPES, pH 7.5, 150 mM NaCl, 0.5 mM TCEP, and 0.2 mg/mL 3xFLAG peptide. The eluted material was subjected to SDS-PAGE and the gel scanned on an Odyssey imager (LI-COR). The gel was then transferred to a nitrocellulose membrane and analyzed by immunoblotting with FLAG antibodies on an Amersham Imager 600 RGB (Cytiva).

For Ufd1-Bpa experiments, Cdc48^E588Q^ (500 nM), HA-tagged, Bpa-incorporated Ufd1 (200 nM), Npl4 (500 nM), and Ub(n)-Dy-sfGFP (1 µM) were mixed in the reaction buffer (50 mM HEPES pH 7.5, 150 mM NaCl, 10 mM MgCl_2_, 0.5 mM TCEP, 0.5 mg/mL protease-free BSA) supplemented with 10 mM of ATP. The reactions were assembled, incubated, and irradiated as described above. After UV irradiation, the protein samples were directly mixed with SDS sample buffer and subjected to SDS-PAGE, followed by silver staining and immunoblotting against the HA epitope.

### Crystallization, data collection, and structure determination

The *ct*UT3-C19 protein (20 mg/mL) was prepared freshly before crystallization setup and centrifuged for 10 min at 15,000 g. Crystals were grown at 20°C in 96-well sitting-drop plates, by mixture of 0.1 μL of protein solution with 0.1 μL of reservoir solution (0.1 M Tris pH 8.0, 28% w/v Polyethylene glycol 4,000) using a protein crystallization robot (Mosquito). Crystals appeared in 7-14 days and were harvested, incubated briefly in cryoprotection solution (0.1 M Tris pH 8.0, 28% w/v Polyethylene glycol 4,000, 10% glycerol), and flash frozen in liquid nitrogen.

All data were collected at BL10U2 beamline of the Shanghai Synchrotron Radiation Facility under 100 K liquid nitrogen stream (wavelength = 0.9798 Å). The crystal contained two copies per asymmetric unit. The *ct*UT3-C19 protein was crystallized in space group C 2 2 21 with cell parameters a = 64.35 Å, b = 67.47 Å, and c = 177.67 Å.

The complex structure was solved by molecular replacement with PHENIX, using the Alphafold2 predicted structure as an initial model. The model was built manually in COOT, refined with PHENIX, and validated using MolProbity. All data processing and structure refinement statistics are summarized in **Table 1**. Structure figures were prepared using PyMOL (http://www.pymol.org/) and ChimeraX.

### Micro-scale thermophoresis (MST)

Synthetic C19 peptide contains an FITC fluorophore. (C19)x(Ub) and (C19scr)x(Ub) conjugates were labeled with fluorescent dye-conjugated maleimide moieties in the HEPES buffer (50 mM HEPES, pH 7.4, 150 mM NaCl, 0.5 mM TCEP) according to the manufacturer’s instructions. Serial dilutions of UT3 or ubiquitin were made by 15 successive 1:1 dilutions from the highest concentration (1560 µM) sample into the HEPES buffer. Each of these solutions was mixed 1:1 with a solution of 2 µM fluorophore-labeled protein or peptide, resulting in the final concentration of the labeled sample being 1 µM and the highest concentration of ligand being 780 µM. All 16 samples were loaded into hydrophilic capillary tubes for 30 min before the final measurements were taken in a Monolith NT.115 instrument (Nanotemper, Munich, Germany). The instrument’s LED (illumination) power was set to 30%, and the MST laser power was set to 40%. Measurements were performed at 25°C. Once the LED was turned on, fluorescence was monitored as a function of time. There was a 5-s waiting period before the MST laser was ignited. The MST laser remained on for 30 s, followed by a 5-s monitoring of the recovery period. The resulting fluorescence time traces were analyzed using MO.Affinity Analysis v2.3 software (NanoTemper Technologies). Averaged relative fluorescence data were fitted in Prism (GraphPad) using the “Dose-response – Stimulation -> [Agonist] vs. response (three parameters)” model algorithm.

### Chemical synthesis of ubiquitin conjugates

The (C19)x(Ub) and (C19scr)x(Ub) conjugates used in this study were synthesized via Fmoc-solid phase peptide synthesis protocols (Fmoc-SPPS) using the standard microwave conditions (CEM Liberty Blue) ^49^. The peptide Ub(C46-K48Ub^Cx19^-C76Acm)-CONH_2_ and Ub(C46-K48Ub^Cx19scr^-C76Acm)-CONH_2_ were synthesized via standard Fmoc-SPPS protocol, Fmoc-Lys(Alloc)-OH and Fmoc-Cys(Acm)-OH were used at Lys48 and Cys76, respectively. The TG XV RAM Rink Amide resin (0.24 mmol/g, Iris Biotech) was used for peptide Ub(C46-K48Ub^Cx19^-C76Acm)-CONH_2_ and Ub(C46-K48Ub^Cx19scr^-C76Acm)-CONH_2_. In brief, 0.4 g of TG XV RAM Rink Amide resin was employed for coupling. 10%(v/v) piperidine containing 0.1 M Oxyma in dimethylformamide (DMF) was used for deprotection condition (1 min at 90°C for each deprotection condition). 0.2 M Fmoc-protected amino acid, 0.14 M N,N’-diisopropylcarbodiimide (DIC) and 0.7 M Oxyma in DMF were used for amino acid coupling conditions at 90°C for 3 min, except for Fmoc-Cys (Acm)-OH, Fmoc-Cys (Trt)-OH, which were coupled at 50°C for 10 min. After the coupling cycles, the resin was incubated with the butoxycarbonyl (Boc) protection mixture containing (di-tertbutyl decarbonate:N,N-diisopropylethylamine:DMF = 1:1:5) for 30 min to introduce the Boc protecting group at the N-terminal of Cys46. The Alloc protecting group was removed by 12 hours incubation (37°C) within 8 mL of dichloromethane (DCM) containing Pd[PPh_3_]_4_ (120 mg) and Ph_3_SiH (400 μL). After incubation, the resin was rinsed by DMF solution containing sodium diethyldithiocarbamate (200 mg/40 mL) to eliminate Pd residual.

Subsequently, the ε-amino group on position Lys48 was further coupled with successive sequence (Ub^Cx19^and Ub^Cx19scr^). After the coupling cycle, the resins were rinsed by DCM and cleaved in TFA cleavage cocktail (82.5% trifluoroacetic acid, 5% thioanisole, 5% tri-isopropylsilane, 2.5% 1,2-ethanedithiol, 5% ddH2O) for 2 hours at 37 ℃. The Crude peptides were precipitated with cold diethyl ether from TFA cleavage mixture and harvested by centrifugation.

The peptide Ub(1-45)-NHNH_2_ was synthesized via standard Fmoc-SPPS protocol^50^. The hydrazine resin (0.4 mmol/g) was used for peptide Ub(1-45)-NHNH_2_. In brief, 0.4 g of hydrazine resin was employed for coupling. The methods of deprotection, coupling and cleavage were same as the synthesis of peptide Ub(C46-K48Ub^Cx19^-C76Acm)-CONH_2_. After TFA cleavage, the crude peptide was obtained by precipitation and centrifugation.

For the synthesis of ubiquitin conjugates (C19)x(Ub) and (C19scr)x(Ub), the native chemical ligation (NCL), desulfurization and deAcm were used. First, Ub(1-45)-MPAA was formed by the crude peptide of Ub(1-45)-NHNH_2_. The method of thioester generation was programmed as previously reported^51^. The Ub(1-45)-MPAA was formed with adding 50 equiv of 2-(4-sulfanylphenyl) acetic acid (MPAA) and 10 equiv of 2,4-pentanedione into peptide Ub(1-45)-NHNH_2_ at 37°C for 2 hours. Then, Ub(1-45)-MPAA was purified by RP-HPLC. Peptide Ub(1-45)-MPAA (2.4 mM,1.2 equiv.) and N-cysteine peptides Ub(C46-K48Ub^x19^-C76Acm)-CONH_2_ (Ub^Cx19^ and Ub^Cx19scr^) (2 mM,1 equiv.) were dissolved in the ligation buffer (0.1 M phosphate, 6 M Gn·HCl, 5 mM TCEP pH 6.8) at room temperature for 10 hours. The reaction was purified by RP-HPLC. The method of superfast desulfurization was programmed as previously reported^50^. The desulfurization reaction was performed at 37°C for 3 minutes under LED light (3.5 W) with standard LEnVLD condition (1 mg/mL peptide, 40 mM TCEP and 5% Rose Bengal (mol%), pH=6.8). The desulfurization product was analyzed by RP-HPLC and verified by ESI MS. Finally, the pH of the solution was then adjusted to 7.3, and Acm protecting group was removed by the addition of PdCl2 (40 mM) at 37°C for 6 hours. The final reaction product was analyzed and purified by RP-HPLC after adding excessive dithiothreitol (DTT) to quench the reaction.

### Liquid chromatography and tandem MS of ubiquitin conjugates

For the purification and characterization of synthetic peptides, reversed-phase high-performance liquid chromatography (RP-HPLC) and LC-ESI-MS were used. The crude peptide Ub(C46-K48Ub^x19^-C76Acm)-CONH_2_ (Ub^Cx19^ and Ub^Cx19scr^) were dissolved in 6 M Gn·HCl containing 0.1 M phosphate (pH 7.5) and purified by RP-HPLC Ultimate XB-C4 (5 μm, 300 Å, Welch) via 20-70% gradient (CH_3_CN/H2O, 1‰ TFA (v/v)) within 30 minutes and characterized via LC-ESI-MS. The crude peptide Ub(1-45)-MPAA was purified by Ultimate XB-C4 (5 μm, 300 Å, Welch) in via 10-60% gradient (CH_3_CN/H2O, 1‰ TFA (v/v)) within 30 minutes and characterized via LC-ESI-MS. The products of NCL were purified by Ultimate XB-C4 (5 μm, 300 Å, Welch) in via 20-70% gradient (CH_3_CN/H2O, 1‰ TFA (v/v)) within 50 minutes and characterized via LC-ESI-MS. The products of desulfrization were purified by Ultimate XB-C4 (5 μm, 300 Å, Welch) in via 10-90% gradient (CH_3_CN/H2O, 1‰ TFA (v/v)) within 30 minutes and characterized via LC-ESI-MS. The final products of ubiquitin conjugates (C19)x(Ub) and (C19scr)x(Ub) were purified by Ultimate XB-C4 (5 μm, 300 Å, Welch) in via 20-70% gradient (CH_3_CN/H2O, 1‰ TFA (v/v)) within 50 minutes and characterized via LC-ESI-MS.

### Band intensity quantifications of gel images

Quantifications of fluorescence scanning gels were carried out using the ImageStudio software (LI-COR). For each lane, a rectangle box was selected to determine total intensity of a band or smear of signal. The rectangle boxes for all lanes on the same gel were kept with similar box sizes. For each gel, an additional rectangle box with similar box size were drawn over an empty or non-signal region to determine background intensity. Signal intensity of each lane was calculated as (total signal intensity – signal box size * background intensity / background box size). The resulting signal intensity was normalized to a designated lane to calculate the relative signal intensity.

## SUPPLEMENTAL INFORMATION

Supplemental information includes:

**Supplemental Document 1**. X-ray Structure Validation Report

**Supplemental Video 1**. Ubiquitin binding to UT3 cleft induces ubiquitin unfolding

## Extended Data Figure Legends

**Extended Data Fig. 1.**
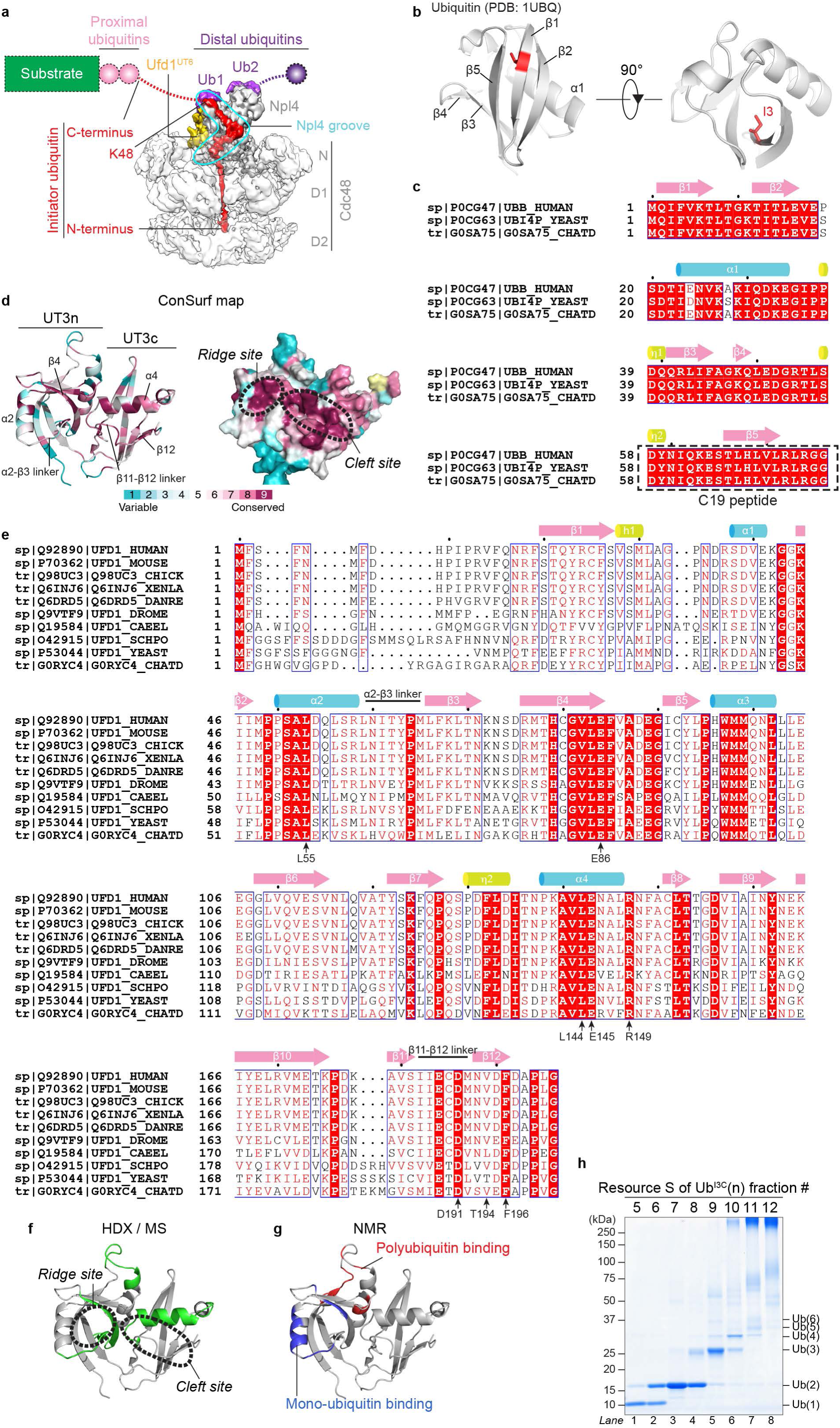
The UT3 domain of Ufd1 contains two highly conserved sites. **a,** Cryo-EM reconstruction of the initiation complex of Cdc48 ATPase (EMD-0665)^15^. Cdc48 is shown in light grey, with the N, D1, and D2 domains indicated. The Npl4 cofactor is colored in dark grey, a C-terminal segment of Ufd1 in golden, the N-terminal segment of initiator ubiquitin in red, and two folded ubiquitin molecules (Ub1 and Ub2) in purple. The “L-shape” Npl4 groove is encircled by a cyan shape. The C-terminal segment of the initiator ubiquitin, proximal ubiquitins, additional distal ubiquitins, and the attached substrate are invisible in the cryo-EM reconstruction and therefore indicated by dashed lines. **b,** Ubiquitin I3 residue is highlighted as red stick in folded ubiquitin (PDB: 1UBQ). Secondary structures are labeled accordingly. **c,** Sequence alignment of ubiquitin proteins from human, *Saccharomyces cerevisiae*, and *Chaetomium thermophilum*. The dashed box highlights the C19 peptides that are identical among the three species. Sequence alignment was generated by Clustal Omega^52^ and visualized by ESPript 3.2^53^. **d,** A conservation map of UT3 was generated using the ConSurf Colab tool and the available UT3 structure (PDB: 1ZC1). Critical secondary structures are labeled in cartoon presentation (*left*). The conserved ridge and cleft sites are highlighted by dashed circles in surface presentation (*right*). The conservation score scale and color coding are shown at the bottom. **e,** Sequence alignment of UT3 from the indicated species was generated and presented as in (**c**). Arrows indicate conserved residues of *Saccharomyces cerevisiae* Ufd1. **f,g,** Shown on the UT3 structure (PDB: 1ZC1) are Ufd1 peptides that previously showed slower HDX upon the binding of polyubiquitinated substrates (green)^15^ (**f**) and residues that showed significant NMR chemical shifts upon the addition of mono-ubiquitin (blue) or polyubiquitin (red)^20^ (**g**). **h,** I3C ubiquitin was used to generate K48-linked polyubiquitin chain and then fractionated via a Resource S column. Proteins in the indicated fractions were analyzed by Coomassie blue staining.

**Extended Data Fig. 2.**
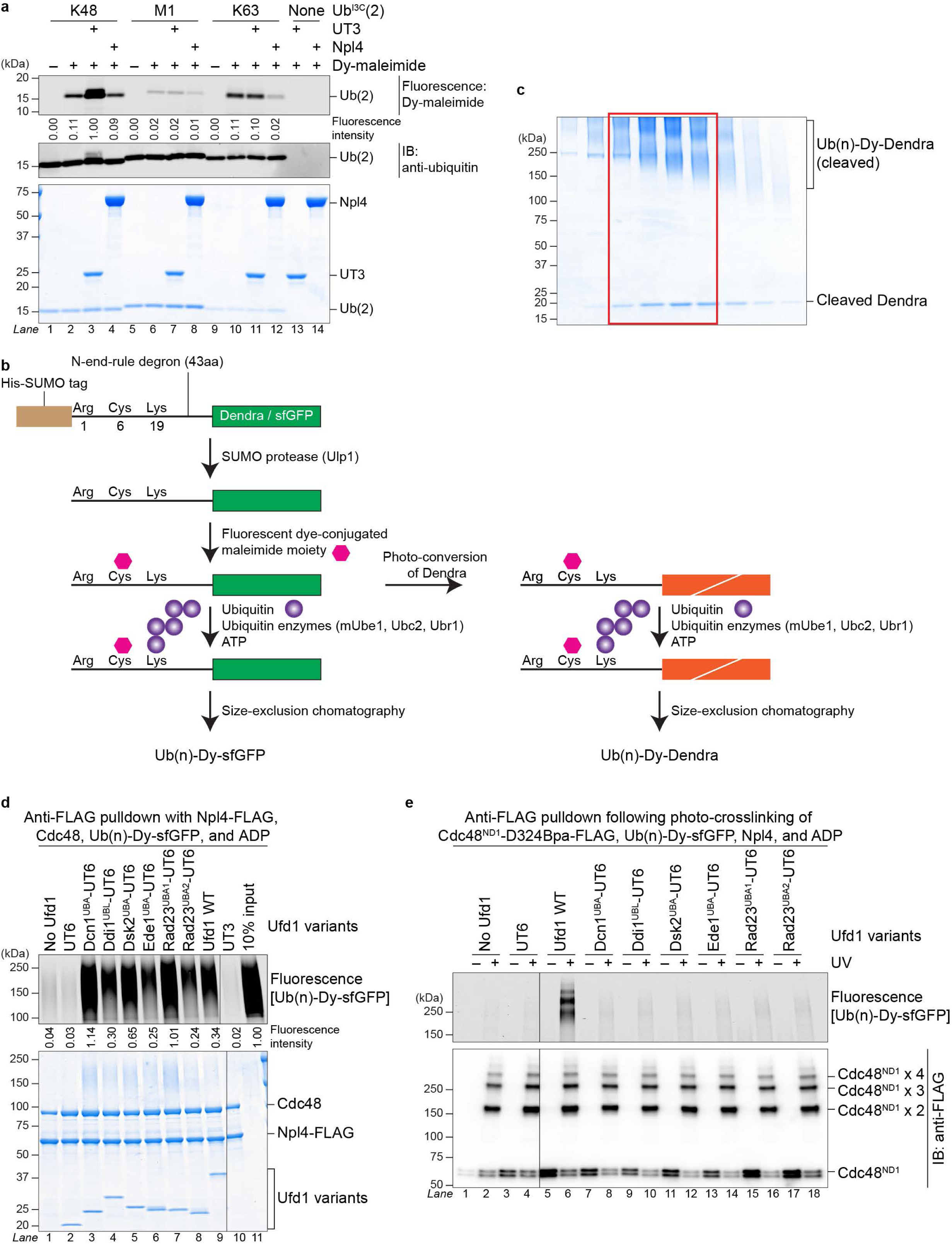
UT3 has a specificity toward K48-linked polyubiquitin and cannot be replaced by other ubiquitin-binding domains. **a,** K48-, M1-, and K63-linked I3C di-ubiquitins were preincubated with UT3 or Npl4 and tested for dye accessibility. Protein mixtures were then analyzed by dye fluorescence scanning (top), anti-ubiquitin blot (middle), and Coomassie blue staining (bottom). **b,c,** A schematic cartoon for the generation of K48-linked polyubiquitinated model substrates containing either Dendra or sfGFP (**b**). The model substrates were covalently labeled with a fluorescent dye, photo-converted by UV as appropriate, and then ubiquitinated, followed by size-exclusion chromatography. A typical gel image of gel filtration fractions is shown (**c**). The red rectangle highlights the pooled fractions. **d,** Polyubiquitinated, dye-labeled sfGFP substrates [Ub(n)-Dy-sfGFP] were mixed with Npl4-FLAG, Cdc48, indicated Ufd1 variants, and ADP, followed by anti-FLAG pulldown. Beads-bound proteins were analyzed by dye fluorescence scanning (top) and Coomassie blue staining (bottom). Dye fluorescence intensities of beads-bound substrates were normalized to input. **e,** Ub(n)-Dy-sfGFP substrates were incubated with indicated Ufd1 variants, Npl4, and a Bpa-incorporated, truncated Cdc48 containing only the N and D1 domains (Cdc48^ND1^). After UV irradiation, Cdc48^ND1^ and proteins covalently crosslinked to it were isolated by anti-FLAG beads. Beads-bound proteins were analyzed by fluorescence scanning (top) and anti-FLAG blot (bottom). The ladder-like band patterns in the immunoblot indicate crosslinks containing multiple molecules of Cdc48^ND1^.

**Extended Data Fig. 3.**
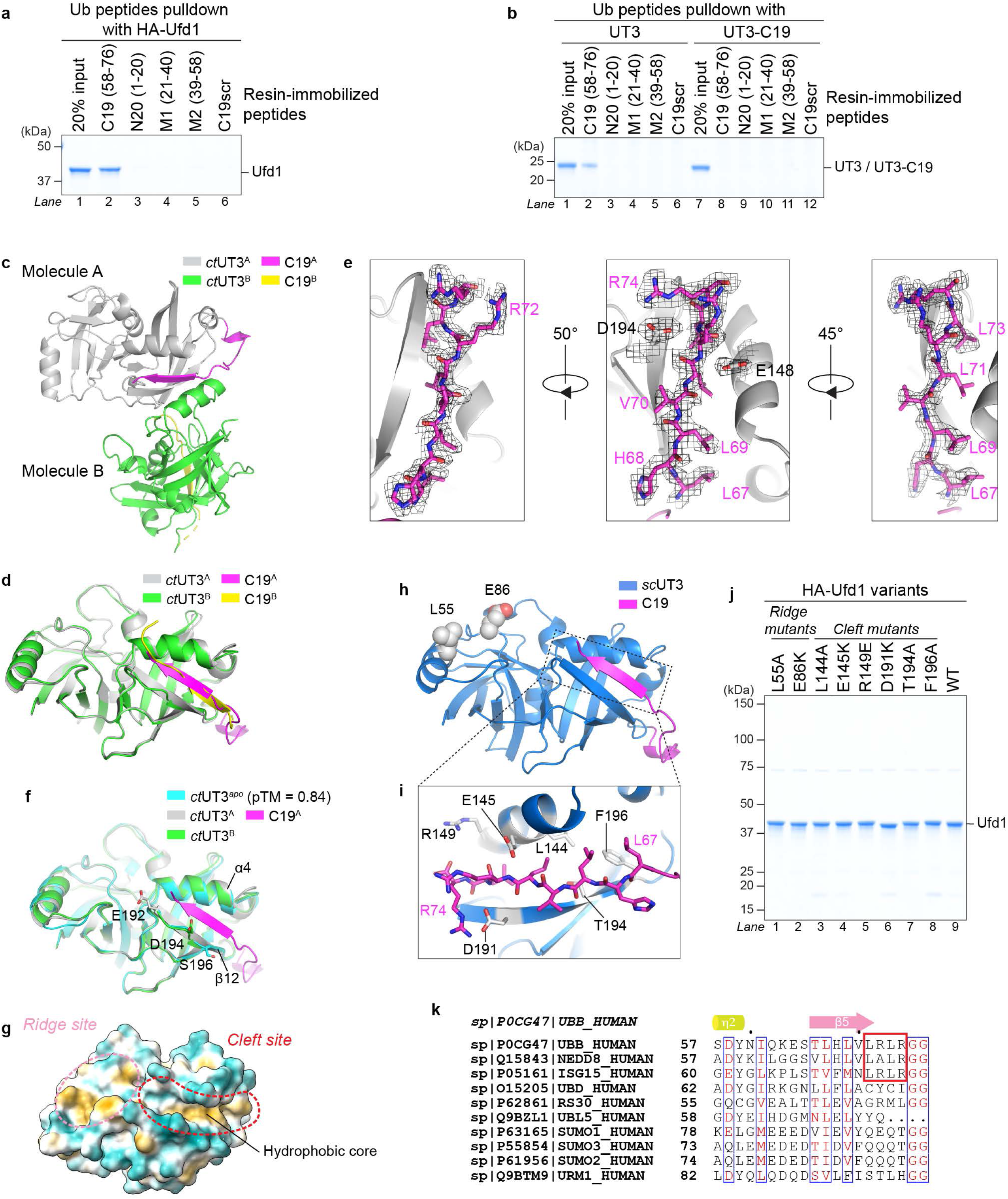
A C-terminal segment of ubiquitin binds to UT3 cleft. **a,b,** Synthetic peptides containing indicated ubiquitin fragments were immobilized and applied for pulldown with full-length Ufd1 (**a**), or UT3 and the UT3-C19 fusion (**b**). Bead-associated proteins were analyzed by Coomassie blue staining. **c,d,** X-ray structure of the *ct*UT3-C19 fusion protein. One asymmetric unit contains two molecules (**c**). Structures of the two molecules are superimposed in (**d**). **e,** Close-up views of the *ct*UT3-C19 structure at the cleft site. *ct*UT3 is shown in grey cartoon and ubiquitin C19 in magenta sticks, overlaid with the 2*F*_o_ − *F*_c_ map at 1.0σ. Residues are numbered with respect to corresponding wild-type proteins. **f,** An AlphaFold3 model of the *apo*-form *ct*UT3 is superimposed to the X-ray structures of *ct*UT3-C19 (molecules A and B). Note that *ct*UT3 β12 starts from S196 in the *apo* model, from D194 in molecule B, and from E192 in molecule A. **g,** Crystal structure of *ct*UT3-C19^A^ is shown with hydrophobicity surface. The C19 peptide at the cleft was removed to reveal the hydrophobic core. **h,i,** A homology model of the *sc*UT3-C19 fusion was generated based on the crystal structure of *ct*UT3-C19^A^. *sc*UT3 is depicted in blue and C19 in magenta (**h**). A close-up view shows the cleft site (**i**). Critical ubiquitin and UT3 residues are highlighted in spheres or sticks. **j,** Wild-type (WT) Ufd1 and indicated Ufd1 variants were expressed recombinantly in bacteria and purified with similar impurities. **k,** Alignment of C19-homology sequences from ubiquitin and ubiquitin-like proteins in human.

**Extended Data Fig. 4.**
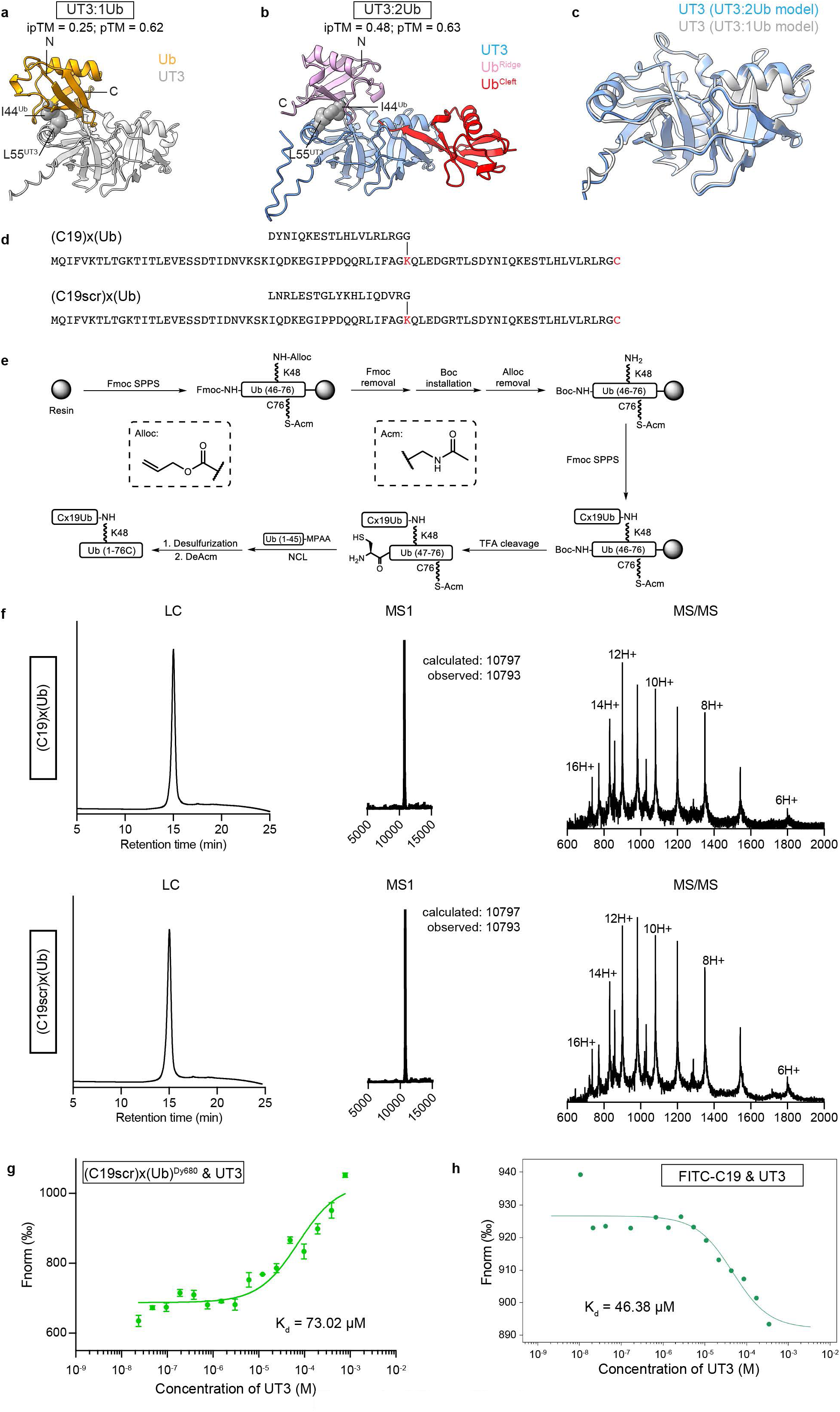
Ubiquitin binding to UT3 ridge and cleft, and chemical synthesis of K48-linked ubiquitin conjugates. **a-c,** AlphaFold3 models of UT3 in complex with one ubiquitin (**a**; UT3:1Ub) and two ubiquitins (**b**; UT3:2Ub) are compared. N and C termini of the ridge-bound ubiquitins are labeled. UT3 L55 and ubiquitin I44 at the interfaces are shown in spheres. The UT3 structures in the two models are superimposed in (**c**). **d,** Primary sequences of the synthesized (C19)x(Ub) and (C19scr)x(Ub) conjugates. Lines between the glycine and lysine residues indicate isopeptide bonds. G76 of the full-length ubiquitin was replaced by a cysteine to allow covalent labeling with a fluorescent dye. **e,** Synthesis roadmap for the ubiquitin conjugates in (**d**). **f,** The (C19)x(Ub) and (C19scr)x(Ub) conjugates were validated by liquid chromatography coupled with tandem MS (LC/MS). **g,h,** Various doses of UT3 were mixed with DyLight 680-labeled (C19scr)x(Ub) conjugates (**g**) or FITC-labeled C19 peptides (**h**), followed by micro-scale thermophoresis measurements. Measurements with (C19scr)x(Ub) were performed in three independent experiments. The data were fitted with non-linear regression to calculate the dissociation constants (K_d_).

**Extended Data Fig. 5.**
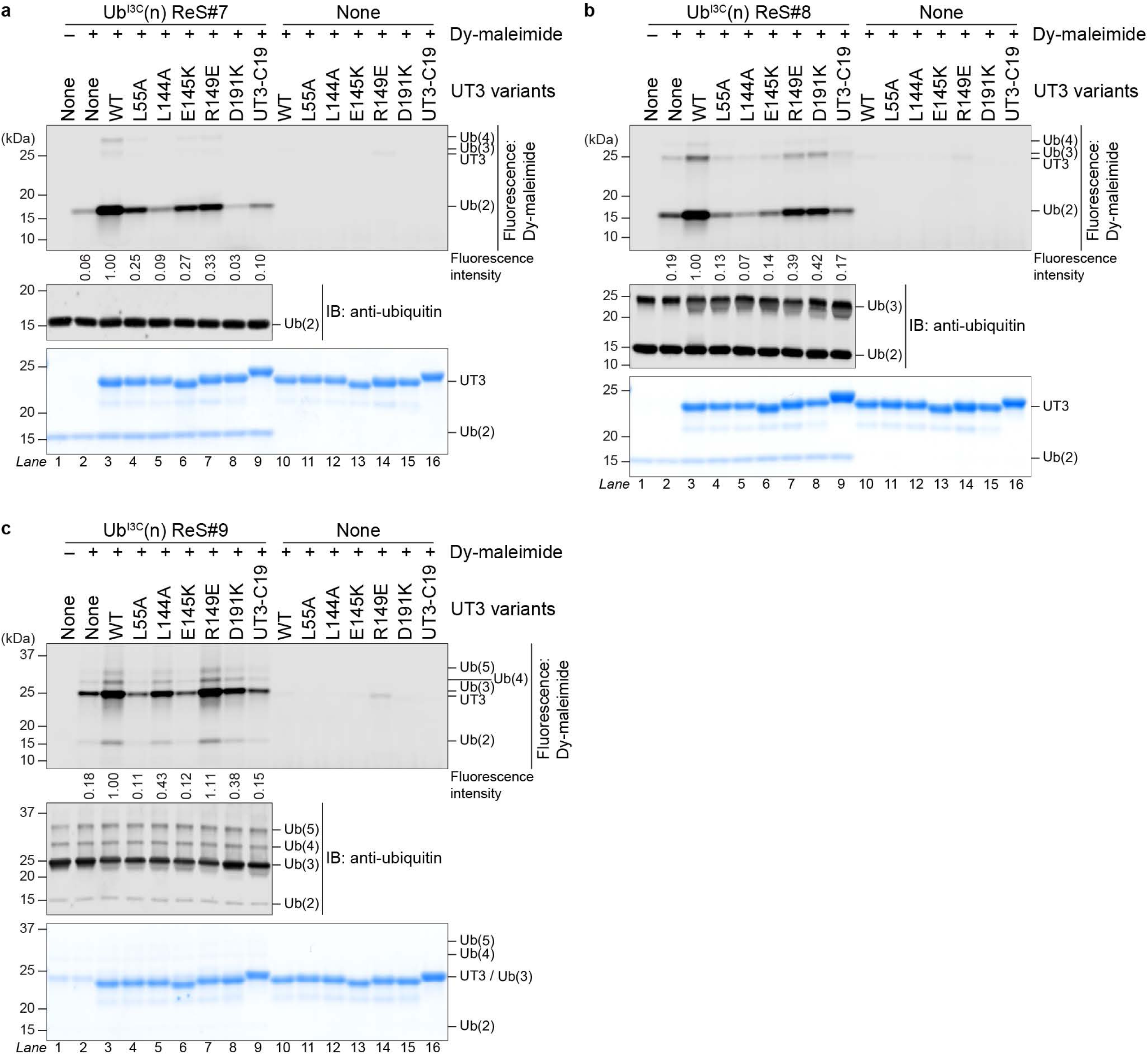
Both the ridge and cleft sites are required for UT3 to unfold ubiquitin. **a-c,** Indicated UT3 variants were tested in dye accessibility assays, with three different fractions of I3C polyubiquitin containing primarily di- and tri-ubiquitin. Protein samples were analyzed by dye fluorescence scanning (top), anti-ubiquitin blot (middle), and Coomassie blue staining (bottom). Intensities of dye fluorescence on di- or tri-ubiquitin were normalized to the corresponding WT UT3 sample.

**Extended Data Fig. 6.**
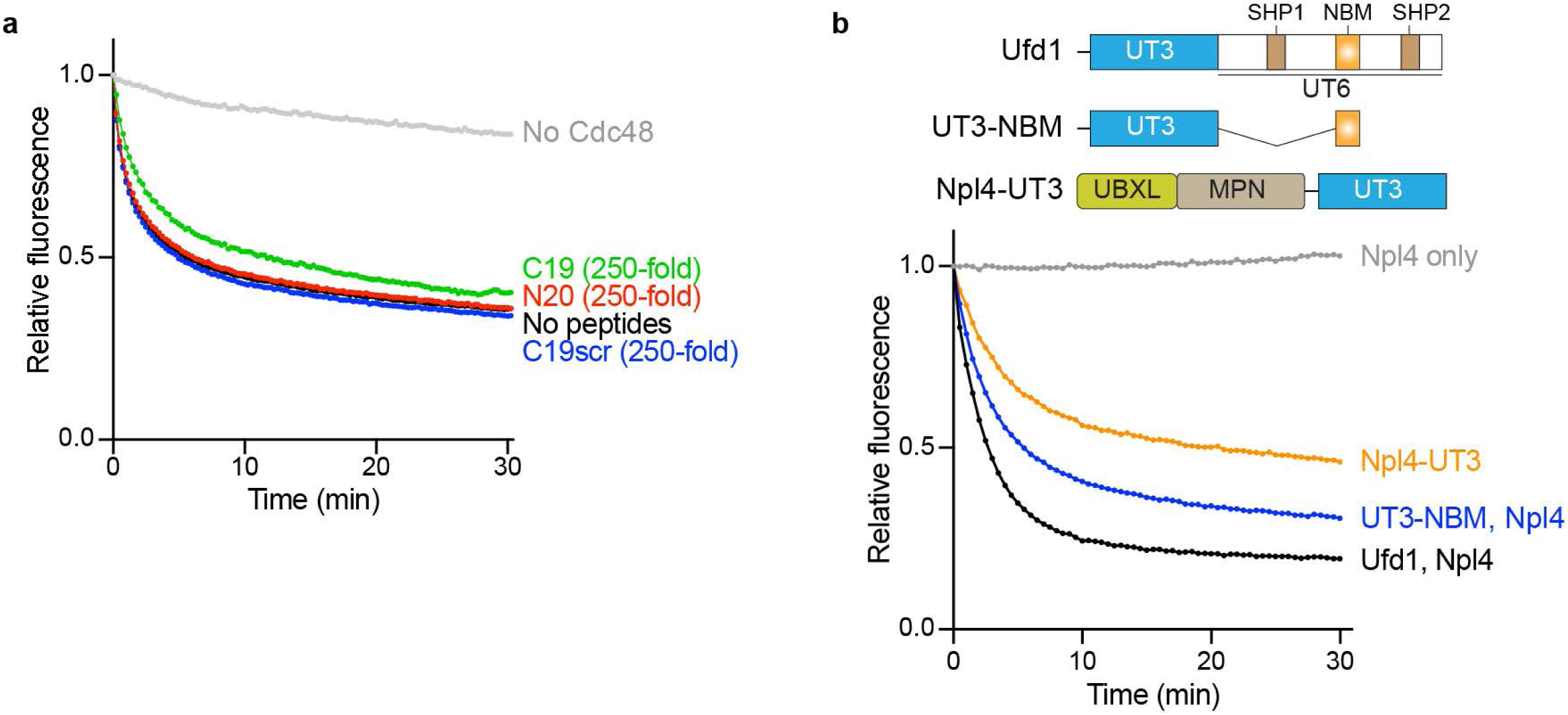
Effects of untethered C19 peptides and variously anchored UT3 on substrate unfolding activity of the Cdc48 ATPase complex. **a,** Polyubiquitinated, photo-converted Dendra substrate was incubated with Ufd1, Npl4, and Cdc48. Synthetic ubiquitin peptides were added in excess as indicated. Upon addition of ATP, the red fluorescence of Dendra was followed over a time course of 30 min. Dendra fluorescence intensities were normalized to the zero-timepoint value of each reaction. **b,** The Dendra substrate unfolding assays were carried out similar to that in (**a**), with the truncated Ufd1 (UT3-NBM) or the Npl4-UT3 fusion. In these cases, most elements of the UT6 domain were omitted.

